# Gram-positive bacterial cell wall components inhibit herpes simplex virus infection

**DOI:** 10.1101/2025.11.02.686169

**Authors:** Amanda N. D. Adams, Lauren E. Griffin, Jonathan Burnie, Jennifer Powers, Amerria Causey, Virginia J. Glick, Miranda Gavitt, Desmond Richmond-Buccola, Cecilia Kim, Maryam Ahmad, Maya Jackson, Griffin Keiser, Jie Lun Cheng, Ambrish Kumar, Liyanage D. Fernando, Jiri Vlach, Megan H. Orzalli, Kizzmekia S. Corbett-Helaire, Alexiane Decout, Parastoo Azadi, Smita Gopinath

## Abstract

The role of the mucosal microbiome in viral infections remains unclear. Genital herpes, caused by herpes simplex virus 1 and 2 (HSV-1 and HSV-2), is among the most prevalent sexually transmitted infections worldwide. Despite evidence linking vaginal *Lactobacillus* to protection against sexually transmitted viruses, the specific microbial components and mechanisms that mediate this defense are not well understood. Here, we show that multiple cell wall components from diverse gram-positive bacteria, including lactobacilli, inhibit HSV-1 and HSV-2 infection in cells and in a mouse model of genital herpes infection. Peptidoglycan (PG) and lipoteichoic acid (LTA), both major components of the gram-positive bacterial cell wall, significantly reduced HSV infectivity *in vitro* and improved survival and disease outcomes in mice. We further showed that *Lactobacillus crispatus* surface layer proteins SlpA and SlpB bind HSV-1 and inhibit infection. Antiviral effects of cell wall components were dose-dependent, relied on intact PG structure, and, in the case of PG and LTA, were independent of TLR2-mediated host signaling. Collectively, our findings identify a species-independent antiviral function for gram-positive bacterial cell wall components against HSV and suggest that the composition of the mucosal microbiome may play an underappreciated role in suppressing mucosal herpes infection in humans.

## Introduction

Genital herpes infection is a chronic, life-long, sexually-transmitted infection that can be caused by both herpes simplex viruses HSV-1 and HSV-2, affecting over 400 million people worldwide (Looker et al., 2015b, 2008). Infected individuals experience recurrent episodes of inflammation and ulceration. Nucleoside analogues like acyclovir that target the viral DNA polymerase are often used to reduce the severity of symptoms once they are detected, but not everyone responds to these medications, and some develop resistance (Schalkwijk et al., 2022). Additionally, symptom recurrence rates are highly heterogenous with some individuals experiencing inflammation and painful ulcers associated with an increased risk of disease transmission, while others display low-level, asymptomatic viral shedding (Casto et al., 2024; Dhankani et al., 2014; Tronstein et al., 2011; Roychoudhury et al., 2020). In addition to transmission between sexual partners, maternal transmission of HSV to neonates, which mostly occurs during birth, can result in 30% fatality rates, even with treatment (Corey and Wald, 2009; De Rose et al., 2023). Thus, there is an urgent need to develop better interventions to block HSV transmission and symptom development.

Mucosal viruses interact with the host microbiome before infecting and replicating within the host. These interactions can influence infectivity, replication, and transmission but have primarily been studied in the mammalian gut mucosa (Pfeiffer and Virgin, 2016). The role of these interactions in the vaginal mucosa remains unclear. The vaginal microbiome of people around the world primarily falls into two categories – a low diversity bacterial community where gram-positive *Lactobacillus* species, like *Lactobacillus crispatus*, dominate, or, a high diversity, gram-negative-dominant, community with low to undetectable *Lactobacillus* (France et al., 2022). Although sometimes found in healthy people (Ravel et al., 2011), the high diversity, *Lactobacillus*-low communities are correlated with an increased risk for bacterial vaginosis (BV), a condition characterized by inflammation, discharge, and pruritus (Abbe and Mitchell, 2023). Clinical studies have shown significant associations between BV and sexually transmitted viral infections including human immunodeficiency virus HIV (Atashili et al., 2008; Borgdorff et al., 2014) human papilloma virus (Qi et al., 2024), and HSV (Cherpes et al., 2003, 2005), suggesting that the presence of *Lactobacillus* decreases the risk of vaginal viral infections in humans.

The protective role of lactobacilli in HSV infection has been previously explored. Pretreatment of mammalian cell lines with vaginal *L. crispatus* reduced HSV-2 viral titers (Mousavi et al., 2018). Furthermore, mice orally colonized with *Lacticaesibacillus rhamnosus* were more protected against vaginal HSV-2 infection, indicating that both gut and vaginal lactobacilli may play a role in influencing HSV-2 risk (Wang et al., 2023). Lactobacilli have also been shown to exhibit antiviral properties via secretion of soluble products including lactic acid and hydrogen peroxide (Conti et al., 2009). Interestingly, the cell-wall of the oral probiotic lactobacillus *Levilactobacillus brevis* was shown to have anti-HSV-2 activity, but the specific antiviral components were not identified (Mastromarino et al., 2011). These studies suggest that while multiple lactobacilli strains have antiviral properties, the specific mechanisms by which they reduce HSV-2 infectivity remain unclear. It also remains unknown if this antiviral protection is specific to probiotic commensals like lactobacilli.

In this study, we sought to identify factors in vaginal bacterial communities that lower HSV-2 risk. By studying health-promoting gut and vaginal lactobacilli, we identify multiple lactobacilli cell wall components including peptidoglycan (PG), lipoteichoic acid (LTA), and S-layer proteins that all inhibit HSV infection. We provide evidence that treatment with these bacterial cell wall components results in little to no inflammation and significantly prolongs survival of infected mice. This antiviral activity of bacterial cell wall components is independent of TLR-2 signaling. Further, we find that this protection extends to PG and LTA isolated from other gram-positive bacteria suggesting that microbial communities rich in gram-positive bacteria play a conserved, protective role against HSV in mucosal environments.

## Results

### *Lactobacillus* cells and isolated peptidoglycan inhibit HSV-2 infection

To determine the mechanisms by which vaginal lactobacilli impact HSV-2 risk, we used *L. crispatus* as a model. To distinguish the impact of the bacterial cell body versus secreted factors on HSV-2 infectivity, we incubated live and dead bacteria with HSV-2 (Figure 1A-C). OD-normalized bacterial cultures were pelleted, washed, and incubated with HSV-2 virions at 37 °C. After incubation, the bacteria were removed by centrifugation and the amount of infectious virus in the supernatant quantified via plaque assay and compared against the control virus incubated without bacteria (Figure 1A-C). We employed two methods of killing: UV exposure (Figure 1B) and phenol (Figure 1C). In both cases, live and killed *L. crispatus* reduced the amount of infectious HSV-2 in the supernatant. These data suggest that the cell body of vaginal *L. crispatus* plays a role in reducing HSV-2 infection *in vitro*.

**Figure 1:**
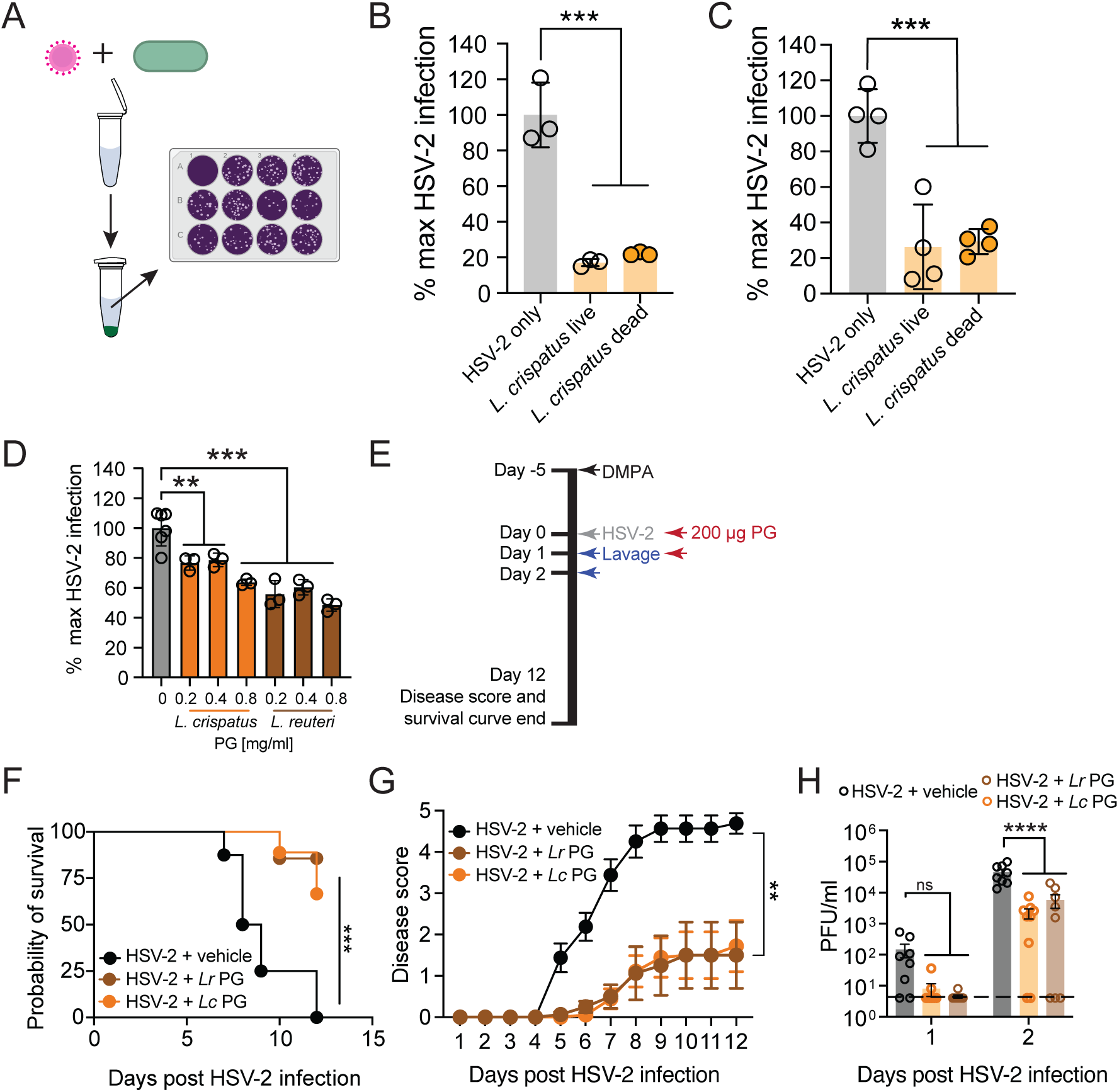
*Lactobacillus* cells and isolated PG inhibit HSV-2 infection. (A) Schematic for the assays shown in (B-C). (B-C) Bacterial cultures of *L. crispatus* MV-1A-US (*Lc*) were grown in NYC III, washed with PBS, OD-normalized to OD_600_ 1.0 and then killed by either UV (B) or phenol (C). (B) A representative experiment is shown n=3, and (C) n=4 across two experiments. Bacteria were then mixed with 40,000 PFU HSV-2 and incubated for one hour before pelleting the bacteria by centrifugation. Infectious HSV-2 in the supernatant was titered by plaque assay on Vero cells as shown in the schematic (A). (B-D) Datapoints are normalized to the average of HSV-2. (D) *L. crispatus* or *L. reuteri* PG (*Lr*) was mixed with 40,000 PFU HSV-2, incubated for two hours, and then the mixture added to Vero cells to quantify the amount of infectious virus using a plaque assay (n=3-6, a representative experiment is shown). (E) Schematic of mouse treatment timeline for experiments shown in (F-H). Mice received 2 mg depot medroxyprogesterone acetate (DMPA) subcutaneously and then five days later were infected intravaginally with 10,000 PFU HSV-2 and either 200 µg *Lc* or *Lr* PG at the time of infection and an additional 200 µg of PG 24 hours later (n=8-9, a representative experiment is shown). Survival was tracked over 12 days (F), and disease severity tracked for 12 days (G). (H) Vaginal lavage was taken for 2 days after the time of infection (starting 24 hours after infection) and lavage viral titers quantified using plaque assays on Vero cells. (H) The dashed line represents the Limit Of Detection (LOD). (B-H) Datasets were compared using one-way ANOVA with Dunnett’s correction (B-D), Log-rank (Mantel-Cox) test (F), and two-way ANOVA with Geisser-Greenhouse correction (G) or Sidak’s correction for multiple comparisons (H). Error bars represent mean and the standard deviation (SD) (B-D) or mean and standard error of the mean (SEM) (G-H).

Like most gram-positive bacteria, most lactobacilli have a thick layer of surface-facing cell wall not enclosed by an outer membrane, consisting largely of PG and teichoic acid (Chapot-Chartier and Kulakauskas, 2014). To identify the components of the *L. crispatus* cell body that influence HSV-2 infectivity, we took a candidate approach and isolated several compounds from the cell wall and determined their impact on HSV-2 infectivity *in vitro* and *in vivo*. We began by purifying PG from *L. crispatus.* Previously, we observed that supernatant from *L. crispatus,* but not human gut-associated *Limosilactobacillus reuteri,* dampened virus-driven inflammation in a mouse model of genital herpes infection (Glick et al., 2024). This suggested that *L. reuteri* was not antiviral against HSV-2 and would make a good negative control. Thus, we also isolated PG from *L. reuteri* for comparison with *L. crispatus* PG in HSV-2 co-incubation assays. Purified *L. crispatus* and *L. reuteri* PG were incubated with HSV-2 for 2 hours at 37 °C before plating the HSV-2-PG mixture on Vero cells to quantify the impact of PG on HSV-2 infectivity via plaque assays (Figure 1D). To our surprise, both *L. crispatus* and *L. reuteri* PG significantly reduced HSV-2 plaques in a dose-dependent manner, with *L. reuteri* PG reducing HSV-2 infectivity slightly better than *L. crispatus*.

Next, we tested if PG could reduce HSV-2 infection *in vivo* using a mouse model of genital herpes infection (Gopinath et al., 2018; Glick et al., 2024; Lebratti et al., 2021; Lee et al., 2020). Conventional C57BL/6 mice were first injected with depot medroxyprogesterone acetate (DMPA) to synchronize estrus cycles and increase sensitivity to HSV-2 infection (Linehan et al., 2004). At the time of infection, we combined 200 μg of *L. crispatus* or *L. reuteri* PG with 10,000 PFU of HSV-2 before infecting mice intravaginally. We also treated mice intravaginally with a second dose of 200 μg of PG on day 1 post-infection (Figure 1E). After vaginal infection in mice, HSV-2 undergoes multiple rounds of local replication in the vaginal mucosa before traveling to the dorsal root ganglia where it continues to replicate before traveling to and infecting fresh epithelial sites resulting in genital inflammation driven by innate immune cells (Lebratti et al., 2021). The virus also spreads to the enteric nervous system and central nervous system resulting in toxic megacolon, paralysis, and eventual death of the host (Khoury-Hanold et al., 2016). To assess the impact of the PG treatment on HSV-2 infection, we tracked disease progression from local genital inflammation to neuronal symptoms over 12 days. We found that mice treated with either *L. crispatus* or *L. reuteri* PG had significantly improved survival (Figure 1F) and significantly lower disease scores when compared to HSV-2 infected control mice (Figure 1G). Vaginal washes were collected for two days after infection to quantify and track infectious virus in the vaginal tract. In this mouse model, early vaginal viral titers are significantly correlated to later stage disease (Shin and Iwasaki, 2012; Gopinath et al., 2018; Lebratti et al., 2021). Two days after infection, the vaginal viral titers were significantly lower in *L. crispatus* and *L. reuteri* PG treated mice than HSV-2 infected control mice (Figure 1H), suggesting that PG blocks HSV-2 early in infection.

### Composition analysis of *L. reuteri* and *L. crispatus* PG

PG structures of food and gut-associated lactobacilli have been previously described (Zhao et al., 2021). However, PG structures of vaginal lactobacilli and *L. reuteri* strain CF48-3A have not been defined. Despite slightly better reduction of HSV-2 infectivity in Vero cells when treated with *L. reuteri* PG as compared to *L. crispatus* PG (Figure 1D), *L. reuteri* and *L. crispatus* PG displayed equivalent protection from HSV-2 disease after intravaginal infection (Figures F-H.) To better understand the differences between our *L. crispatus* and *L. reuteri* PG preparations, we assessed the purity of the isolated PG and further defined its structure using GC-MS of trimethylsilyl (TMS) methyl glycosides to detect *N*-acetylglucosamine (GlcNAc) and *N*-acetylmuramic acid (MurNAc), and GC-MS after acid hydrolysis and derivatization with heptafluorobutyric anhydride (HFBA) to detect amino acids. The results (Table 1) indicate that the purified *L. crispatus* and *L. reuteri* PG samples are enriched in amino acids that are commonly found in PG, including alanine, glutamate, lysine, and glycine. Overall ratios of amino acids were similar between *L. crispatus* and *L. reuteri*, indicating that the amino acid content of the PG is similar between these two strains. Typical PG amino acids are lysine, glycine, glutamate, and alanine (Rohde, 2019). The results also indicated the presence of other amino acids that are not commonly found in PG, including proline, tyrosine and phenylalanine which may be due to contaminating proteins. Glucosamine and muramic acid were present in a 1:1 ratio for both PG samples as would be expected (Table 1).

**Table 1:** *Composition Analysis of *L. reuteri* and *L. crispatus* PG.

| Analysis | Before HF |  | After HF |  |
| --- | --- | --- | --- | --- |
|  | *L. reuteri* | *L. crispatus* | *L. reuteri* | *L. crispatus* |
| **Amino acids (HFBA)** |  |  |  |  |
| Alanine (Ala) | 29.6 | 30.3 | 22.7 | 23.2 |
| Glycine (Gly) | 12.8 | 13.2 | 14.3 | 14.8 |
| Valine (Val) | 4.3 | 4.3 | 5.6 | 4.7 |
| Threonine (Thr) | 3.7 | 3.6 | 4.6 | 4.5 |
| Serine (Ser) | 3.7 | 3.5 | 4.6 | 4.9 |
| Leucine (Leu) | 8.3 | 8.2 | 10.9 | 10.8 |
| Isoleucine (Ile) | 2.6 | 2.6 | 3.4 | 3.3 |
| Proline (Pro) | 11.7 | 11.3 | 10.9 | 11.8 |
| Aspartate (Asp) | 4.6 | 4.6 | 6.5 | 6.6 |
| Glutamate (Glu) | 6.4 | 6.3 | 5.5 | 5.4 |
| Phenylalanine (Phe) | 2.1 | 2.1 | 2.8 | 3.8 |
| Lysine (Lys) | 8.8 | 8.6 | 5.9 | 5.0 |
| Tyrosine (Tyr) | 1.5 | 1.4 | 2.3 | 1.4 |
| Total Amino Acids | 100 | 100 | 100 | 100 |
| **Carbohydrate (TMS)\*\*** |  |  |  |  |
| Glucosamine | 45 | 47 | 46 | 43 |
| Muramic acid | 55 | 53 | 54 | 57 |
| Total Sugars | 100 | 100 | 100 | 100 |

\* Composition is given as relative mole percent. \*\* Besides the two carbohydrate residues, glycerol and amino acids were detected but not quantified.

Besides the two PG carbohydrates, both *L. crispatus* and *L. reuteri* PG isolates contained glycerol that was greatly reduced upon hydrofluoric acid treatment (Supplementary Figure 1), consistent with the presence of Wall Teichoic Acids (WTAs) in the untreated PG. While LTAs are removed by SDS during the isolation, WTA can be expected given its covalent attachment to PG. Composition analysis of the HF-treated PG showed that the relative ratios of amino acids as well as GlcNAc and MurNAc were very similar to the untreated PG (Table 1). Further analysis of the HF-treated PG by ^1^H NMR showed signals mainly due to PG carbohydrates and amino acids, without apparent contaminants (Supplementary Figure 2). Taken together, these results indicate that the PG preparations both likely contain an associated WTA and are low in other co-purifying contaminants. Additionally, the PG amino acid and amino sugar contents of *L. crispatus* and *L. reuteri* are nearly the same, in-line with our observation that these PGs display similar levels of antiviral activity against HSV-2.

### *L. crispatus* PG suppresses HSV-2 infection independently of the host microbiome

Since the mouse vaginal mucosa is host to endogenous gram-positive bacterial species (Vrbanac et al., 2018), we wanted to determine if the presence of the endogenous vaginal microbiome was required for PG-mediated antiviral activity. We infected germ-free C57BL/6 mice intravaginally with HSV-2 with and without 100 μg of *L. crispatus* PG at the time of infection (Supplementary Figure 3). In this experiment, we did not include a second treatment of *L. crispatus* PG on day 2 post-infection to minimize mouse handling that could contaminate the experiment. However, despite only receiving a single dose of PG, we found that *L. crispatus* PG significantly improved survival (Supplementary Figure 3A-B) and symptom development (Supplementary Figure 3C).

This protection was accompanied by a reduction in vaginal viral titers (Supplementary Figure 3D) like that observed in mice with a conventional microbiome, indicating that *L. crispatus* PG protection from HSV-2 does not require the presence of an endogenous microbiome. Our data also suggest that a reduced dose and reduced frequency of PG treatment is still protective.

### Robust *L. crispatus* PG inhibition of HSV-2 infection requires PG at time of infection

Next, to assess whether PG is required only at the time of infection to protect from HSV- 2 disease, we first infected mice intravaginally without PG and then treated them with *L. crispatus* PG 4 hours after infection and once daily for 7 days post-infection (Supplementary Figure 4). Post-infection treatment did not ameliorate disease progression. PG treated mice had equivalent survival outcomes (Supplementary Figure 4A-B) and similar disease progression (Supplementary Figure 4C) as control mice despite a reduction in vaginal viral titers 2 days post-infection (Supplementary Figure 4D). This was in striking contrast to the significant inhibition of disease observed upon addition of PG at the time of infection (Figure 1). This further suggests that *L. crispatus* and *L. reuteri* PG presence is required during early viral infection to restrict HSV-2 inflammation and disease and that the second dose of PG on day 1 post-infection was not required for protection.

### Gram-positive PG reduces HSV-2 infection *in vitro* and *in vivo*

Since both *L. crispatus* and *L. reuteri* PG were equivalently protective of HSV-2 infection *in vivo*, we asked whether this protection was broadly true of PG. We assessed the impact of commercially available PG from *Bacillus subtilis,* a gut commensal, and *Staphylococcus aureus,* a skin commensal, on HSV-2 infection in cells (Figure 2A-B) and in mice (Figure 2C-I). Both *B. subtilis* and *S. aureus* PG significantly reduced HSV- 2 infections in Vero cells at doses lower than those seen with *L. reuteri* and *L. crispatus* PG [0.2 mg/ml] (Figure 1D), with *B. subtilis* PG being more protective than *S. aureus* PG (Figure 2A-B). Based on the increased antiviral efficacy observed in cells, we tested if a smaller dose of 50 µg of PG could protect mice against genital HSV-2 infection. We found 50 µg of *B. subtilis* PG completely rescued HSV-2 infected mice (Figure 2D-F) with PG-treated mice showing no symptoms of infection (Figure 2E) and containing no detectable virus in the vagina (Figure 2F). *S. aureus* PG was also highly protective and rescued survival and disease (Figure 2G-H) with significantly lower viral titers than HSV- 2 infection alone two days post-infection (Figure 2I). To determine if this was unique to gram-positive PG, we also treated mice with the same amount of gram-negative *Escherichia coli* PG at the time of infection (Figure 2J-L). *E. coli* PG did not rescue survival (Figure 2J) or disease symptoms (Figure 2K), and infected mice had equivalent levels of vaginal virus with or without PG treatment (Figure 2L). Taken together, these results demonstrate that gram-positive PG is protective against HSV-2 infection.

**Figure 2:**
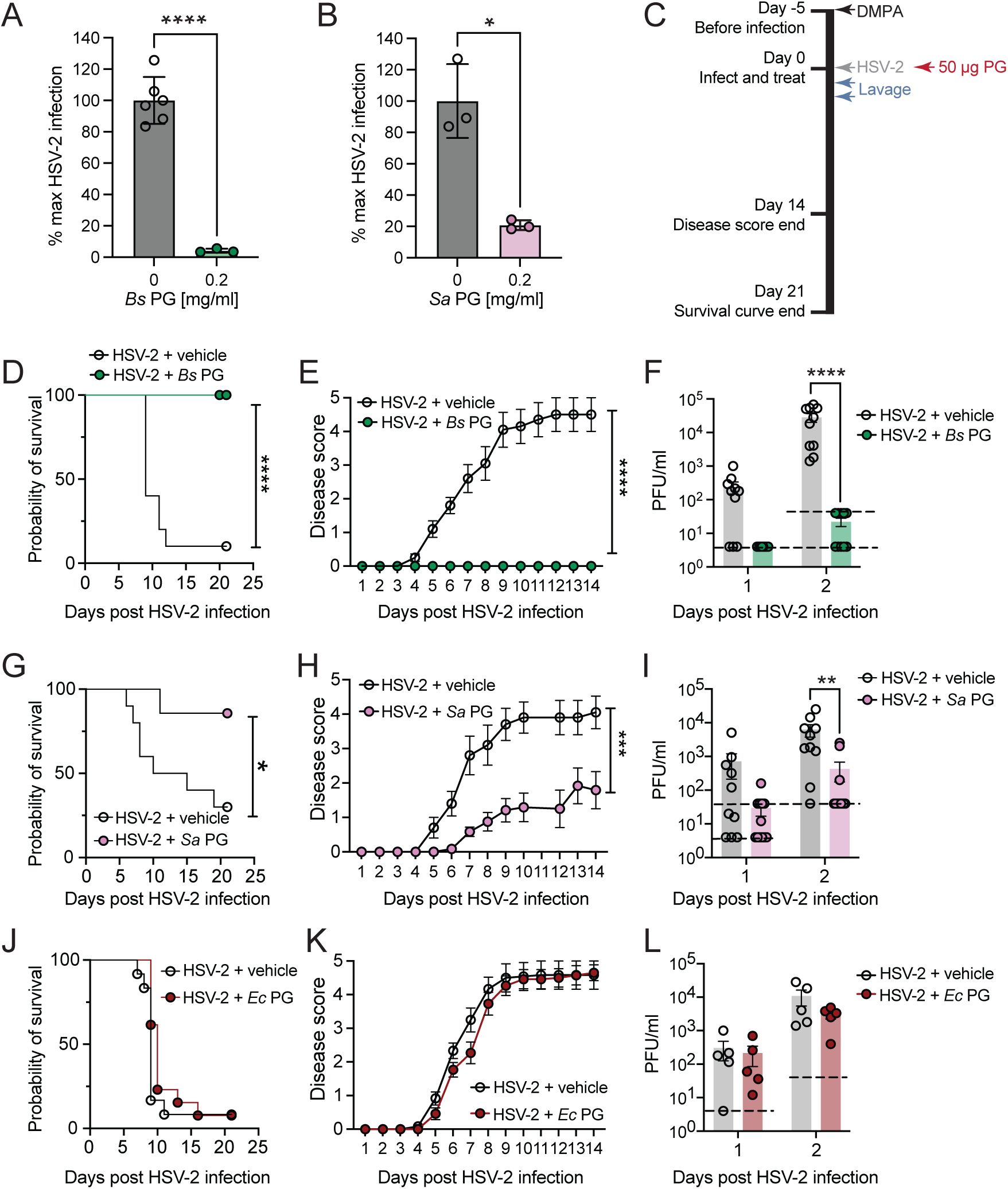
Gram-positive PG reduces HSV-2 infection *in vitro* and *in vivo*. (A-B) *B. subtilis* (*Bs*) and *S. aureus* (*Sa*) PG (both from Sigma-Aldrich) were incubated with 40,000 PFU HSV-2 for two hours and then the amount of infectious virus in the mixture quantified using plaque assays on Vero cells (n=3-6, a representative experiment is shown). (C) Schematic of mouse treatment timeline for the experiments shown in (D-L). Five days after DMPA treatment, mice were infected intravaginally with 10,000 PFU HSV-2 and 50 µg *B. subtilis* PG (Sigma-Aldrich) (D-F), *S. aureus* PG (G-I) (InvivoGen), or *E. coli* PG (*Ec*) (InvivoGen) (J-L). Survival was tracked over 21 days (D, G, and J) and disease scores notes for 14 days (E, H, and J). Vaginal lavage was collected on days 1 and 2 post-infection and infectious virus titered via plaque assay (F, I, and L). (D-F, G-I, and J-K) n=10-12 across two independent experiments. (L) n=5 from a representative experiment. The LOD for (F) on day 2 post-infection was 40 PFU/ml for one experiment (5 mice) and 4 PFU/ml for the second experiment (5 mice). LOD for (I) on day 1 was 4 PFU/ml for one experiment (12 mice) and 40 PFU/ml for the second experiment (10 mice) and on day 2 40 PFU/ml for both experiments (22 mice). LOD for (L) was 4 PFU/ml on day 1 and 40 PFU/ml on day 2. Data were compared using Welch’s t-test (A-B), Log-rank (Mantel-Cox) test (D, G, and J) and two-way ANOVA with Geisser-Greenhouse correction (E, H, and K) or Sidak’s correction (F, I, and K). Error bars represent mean and SD (A-B) or mean and SEM (E, F, H, I, K, and L).

### PG protection of HSV-2 infection is lysozyme sensitive

PG preparations from commercial sources can be complex mixtures with co-purifying components that can impact viruses (Johnson et al., 2019). To assess whether commercial PG protection from HSV infection was dependent on PG rather than co-purifying compounds, we treated *B. subtilis* PG with lysozyme, a muramidase, which preferentially cleaves the disaccharide MurNAc and GlcNAc backbone of PG. Treatment of *B. subtilis* PG with lysozyme increased infectivity of HSV-2 in cells compared to untreated PG (Figure 3A). However, the monosaccharides GlcNAc and MurNAc alone did not reduce HSV-2 infection in cells (Figure 3B). Treatment of *B. subtilis* PG with lysozyme also reduced *B. subtilis* PG protection of HSV-2 in mice, while lysozyme did not have a significant impact on survival, disease, or vaginal viral titers as compared to HSV-2 alone (Figure 3C-E). Lysozyme treatment of *B. subtilis* PG increased the detectable virus in the vagina as compared to untreated *B. subtilis* PG (Figure 3E). This demonstrates that PG protection from HSV-2 is lysozyme sensitive, suggesting a possible role for longer PG linkages in protecting against HSV-2 infection.

**Figure 3:**
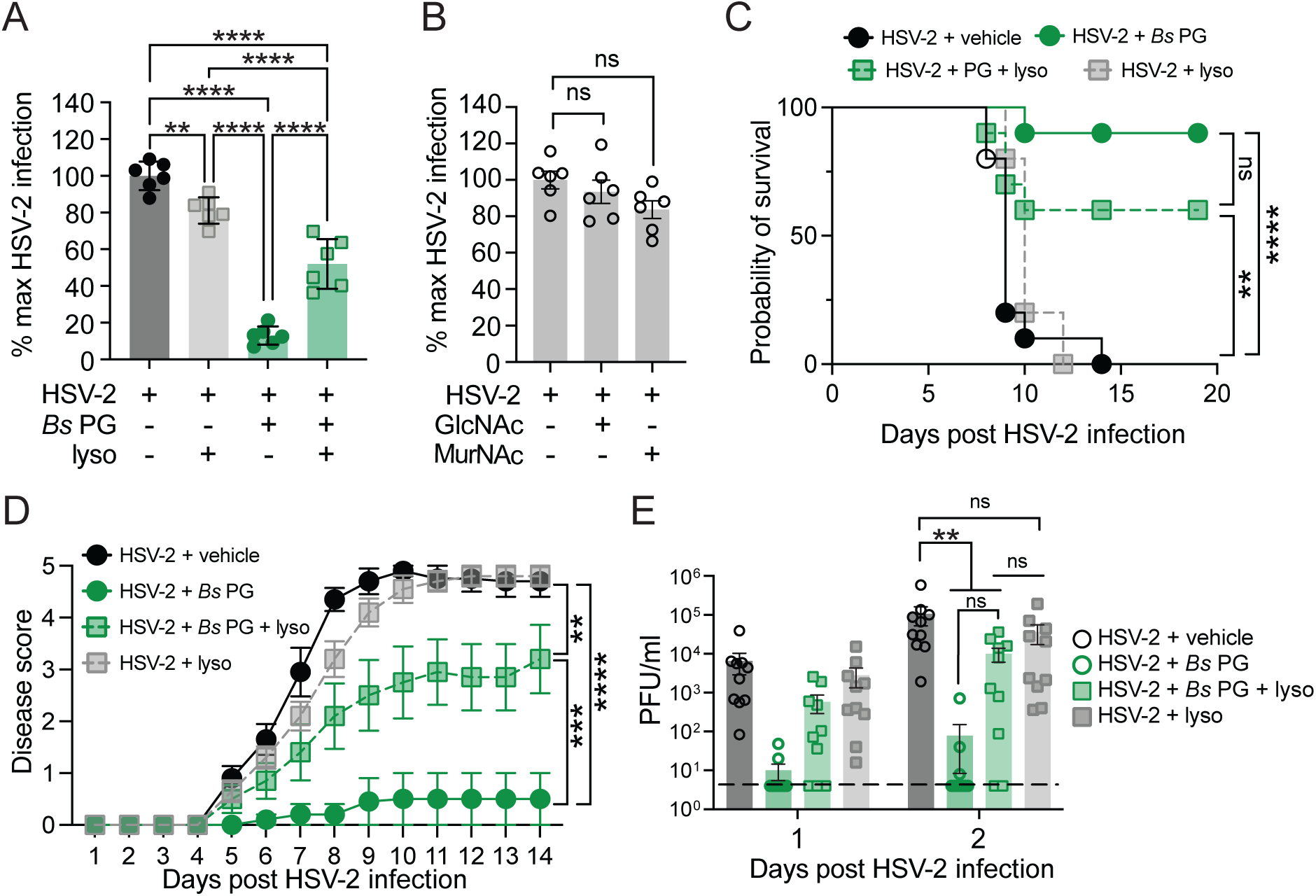
PG protection of HSV-2 infection is lysozyme sensitive. (A) [0.05 mg/ml] *B. subtilis* PG (Sigma-Aldrich) with or without 1 mM lysozyme (lyso) was mixed with 40,000 PFU HSV-2, incubated for 2 hours and then the mixture added to Vero cells to quantify the amount of remaining infectious virus using a plaque assay (n=6, across two independent experiments). (B) [0.2 mg/ml] GlcNAc or MurNAc was mixed with 40,000 PFU HSV-2, incubated for two hours, and then the mixture added to Vero cells to quantify the amount of infectious virus using a plaque assay. (n=6, across two independent experiments). (C-E) Five days post DMPA-treatment, mice (n=10, across two independent experiments) were infected intravaginally with 10,000 PFU HSV-2 and the following treatments at the time of infection: 50 µg of *B. subtilis* PG (Sigma-Aldrich) with or without 10 mM lysozyme or 10 mM lysozyme alone. Survival was tracked over 20 days (C), and disease severity tracked for 14 days (D). (E) Vaginal lavage was collected for two days and infectious virus quantified using plaque assays. Samples were compared using one-way ANOVA with Tukey’s (A) or Dunett’s correction (B), Log-rank (Mantel-Cox) test (C), and two-way ANOVA with Geisser-Greenhouse correction (D) or Tukey’s correction (E). Error bars represent mean and SD (A) or mean and SEM (B, D, and E).

### *B. subtilis* PG inhibits HSV-1 infection in human fibroblasts

Given the robust protection of gram-positive PG on HSV-2 infection, we asked whether PG could also protect against HSV-1 infection. First, using HSV-1 K26GFP where GFP is fused to viral protein VP26 (Desai and Person, 1998), we monitored virion entry and replication of GFP-labelled capsid containing viral particles. In primary human foreskin fibroblasts (HFFs), GFP expression was undetected in cells infected with HSV-1 and *B. subtilis* PG 6 hours post-infection (Supplementary Figure 5A). We observed significantly reduced frequencies of GFP+ infected cells at 24 hours post-infection across a range of viral infectious doses (Supplementary Figure 5B). This protection was also maintained in a second barrier cell type, normal human epidermal keratinocytes (NHEK) (Orzalli et al., 2021) (Supplementary Figure 6). In the presence of [1 mg/ml] *S. aureus* PG, HSV-1 K26GFP infection of NHEK was blocked at multiple infectious doses (MOI 1 and 10) (Supplementary Figure 6A-B and E-F). Lower levels of *S. aureus* PG [0.25 mg/ml] only blocked infection at MOI of 1 (Supplementary Figure 6C-D) but not at MOI of 10 (Supplementary Figure 6G-H). These data show that gram-positive PG blocks both HSV-1 and HSV-2 infection in barrier cells.

### Lipoteichoic acid inhibits HSV-1 and HSV-2 infection

Since teichoic acids are another abundant hallmark of gram-positive bacterial cell walls as compared to gram-negative bacteria, we wanted to test if LTA had an independent impact on antiviral protection. Thus, we sought to purify LTA from *L. crispatus* and *L. reuteri* to determine their impact on HSV infection. We were successfully able to isolate *L. reuteri* LTA but were unable to isolate *L. crispatus* LTA using the same techniques (see Materials and Methods and Supplementary Figure 7). We then tested *L. reuteri* LTA impact on HSV-1 and HSV-2 infection (Figure 4). First, we tested LTA efficacy against HSV-1 infection in human endocervical cells (End1) (Figure 4A-C). *L. reuteri* LTA, as well as commercial *B. subtilis* and *S. aureus* LTA were mixed with HSV-1 and added to cells at an MOI of 1. LTAs from all three bacterial species significantly inhibited HSV-1 infection in End1 cells. Both *L. reuteri* and *B. subtilis* LTA reduced HSV-1 infection rates by 99.9% while *S. aureus* LTA was slightly less suppressive, inhibiting infection by an average of 82.5% ± 1% SD. We also tested the impact of LTA from *B. subtilis*, *S. aureus,* and the gram-positive throat and vaginal commensal *Streptococcus pyogenes* on HSV-1 infection in HFFs (Figure 4D-E). When mixed with virus and added at the time of infection, all three LTAs blocked HSV-1 infection to nearly undetectable levels, with *Bs* LTA reducing the frequency of infected cells from 89% ± 1% SD to 2.3% ± 0.3% SD (Figure 4D-E). We then evaluated the impact of *L. reuteri* LTA on genital HSV-2 infection in mice by combining 50 µg of *L. reuteri* LTA alongside HSV-2 at the time of infection. *L. reuteri* LTA completely blocked HSV-2 disease and we recovered no vaginal viral titers from these mice two days post-infection (Figure 4F-H). Collectively, these data show that, in addition to PG, LTA is a unique feature of gram-positive bacteria that robustly reduces HSV-1 and HSV-2 infection.

**Figure 4:**
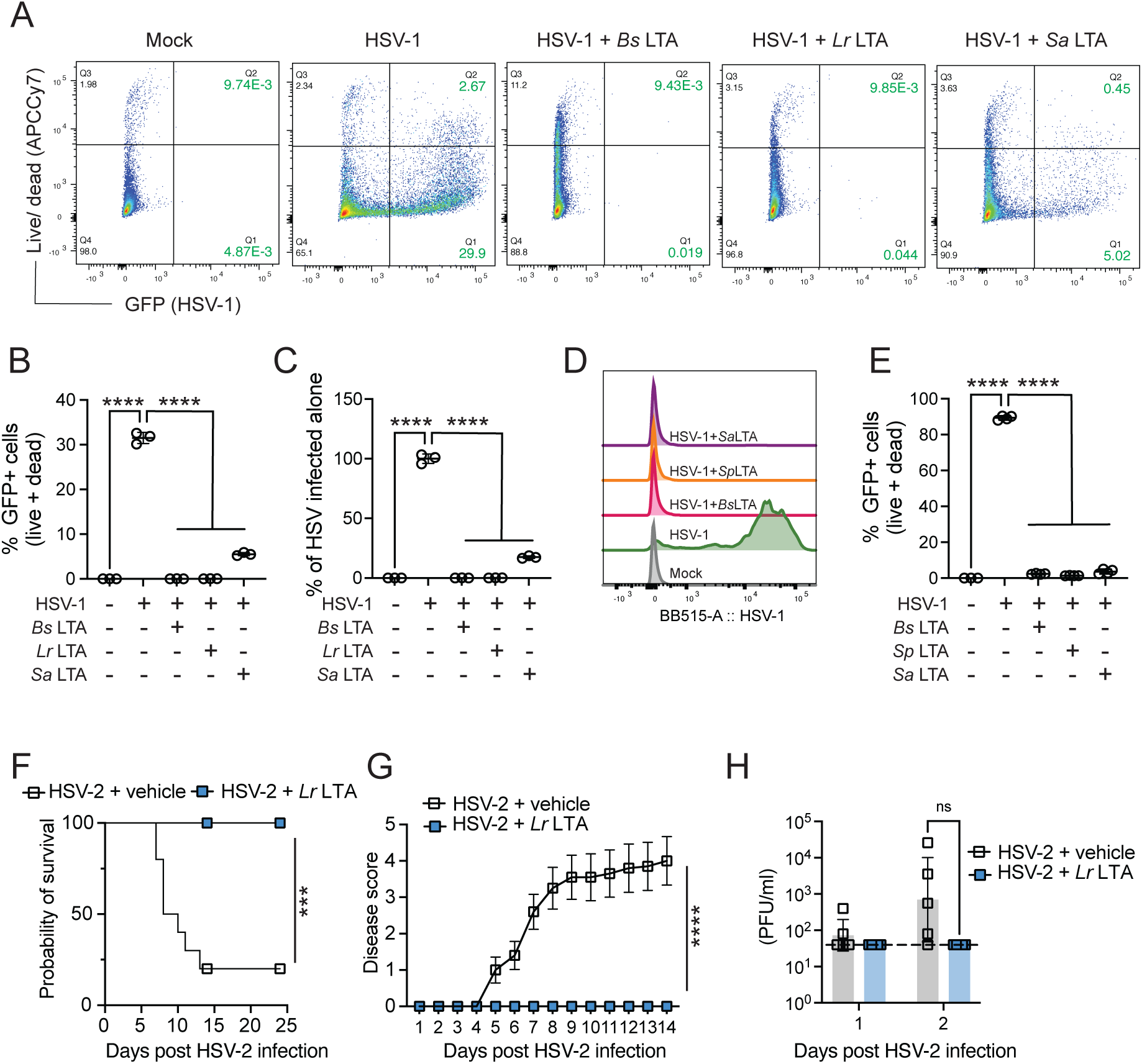
LTA inhibits HSV-1 and HSV-2 infection. 50 μg LTA from *B. subtilis* (InvivoGen)*, L. reuteri* and *S. aureus* (InvivoGen) was incubated with HSV-1 K26GFP (MOI 1) and added to human End1 cells and frequency of live and dead GFP+ cells quantified (A-B) and normalized against HSV-1 infection condition alone (C) 22 hours post-infection (n=3 a representative experiment shown). 50 μg LTA from *B. subtilis, S. pyogenes* (*Sp*) (Sigma-Aldrich), and *S. aureus* was incubated with HSV-1 K26GFP (MOI 1) and added to HFF cells and GFP expression from representative conditions shown (D) and quantified (E) 22 hours post-infection (n=4, a representative experiment is shown). (F-H) Five days after DMPA treatment, mice (n=10, across two independent experiments) were infected with 10,000 PFU HSV-2 alone or with 50 µg *L. reuteri* LTA. Survival was tracked over 20 days (F), and disease severity tracked for 14 days (G). Vaginal lavage was collected and infectious virus quantified using plaque assays (n=5, a representative experiment is shown) (H). P-values were calculated using one-way ANOVA with Dunnett’s correction (B-C, and E), Log-rank (Mantel-Cox) test (F), and two-way ANOVA with Geisser-Greenhouse correction (G) or Sidak’s correction (H). Error bars represent mean and SD (B, C, and E) or mean and SEM (G-H).

### *L. crispatus* surface layer proteins inhibit HSV infections

In addition to the conserved PG and LTAs expected of all gram-positive bacteria, multiple members of lactobacilli species have surface layer proteins that form a 2- dimensional lattice-like layer covalently linked to cell wall components (Palomino et al., 2023). Recent work has identified the presence of two surface layer proteins, Slp A and SlpB, in vaginal *L. crispatus* species (Decout et al., 2024). To test if *L. crispatus* SLPs could inhibit HSV-1 infection of HFFs, we combined SlpA or SlpB protein with HSV-1 at the time of infection at an MOI of 1 (Figure 5A-B) and MOI of 10 (Figure 5C-E). We found 8 μg [0.032 mg/ml] of SlpA or SlpB was sufficient to significantly inhibit HSV-1 infection, reducing infection rates from 83% ± 6% SD to fewer than 1% of all cells at an MOI of 1 (Figure 5A-B). This inhibition was maintained but significantly reduced with a higher infectious dose (Figure 5C-E). At an MOI of 10, higher concentrations of the protein only reduced infection by an average of 15% (SlpA) and 17% (SlpB). We observed a three-fold decrease in GFP mean fluorescence intensity indicating significant restriction of viral replication (Figure 5D). While lower doses of 1 µg of SlpA or SlpB proteins failed to protect cells at 22 hours post-infection (Figure 5C), we observed significant reduction of the frequency of infected cells at 6 hours post-infection (Figure 5E) suggesting that this inhibition can be overcome at lower doses of SlpA/SlpB or higher viral infectious doses. These data suggest that in addition to PG and LTA, the outermost protein layers of *L. crispatus* can significantly inhibit HSV-1 infection in cells.

**Figure 5:**
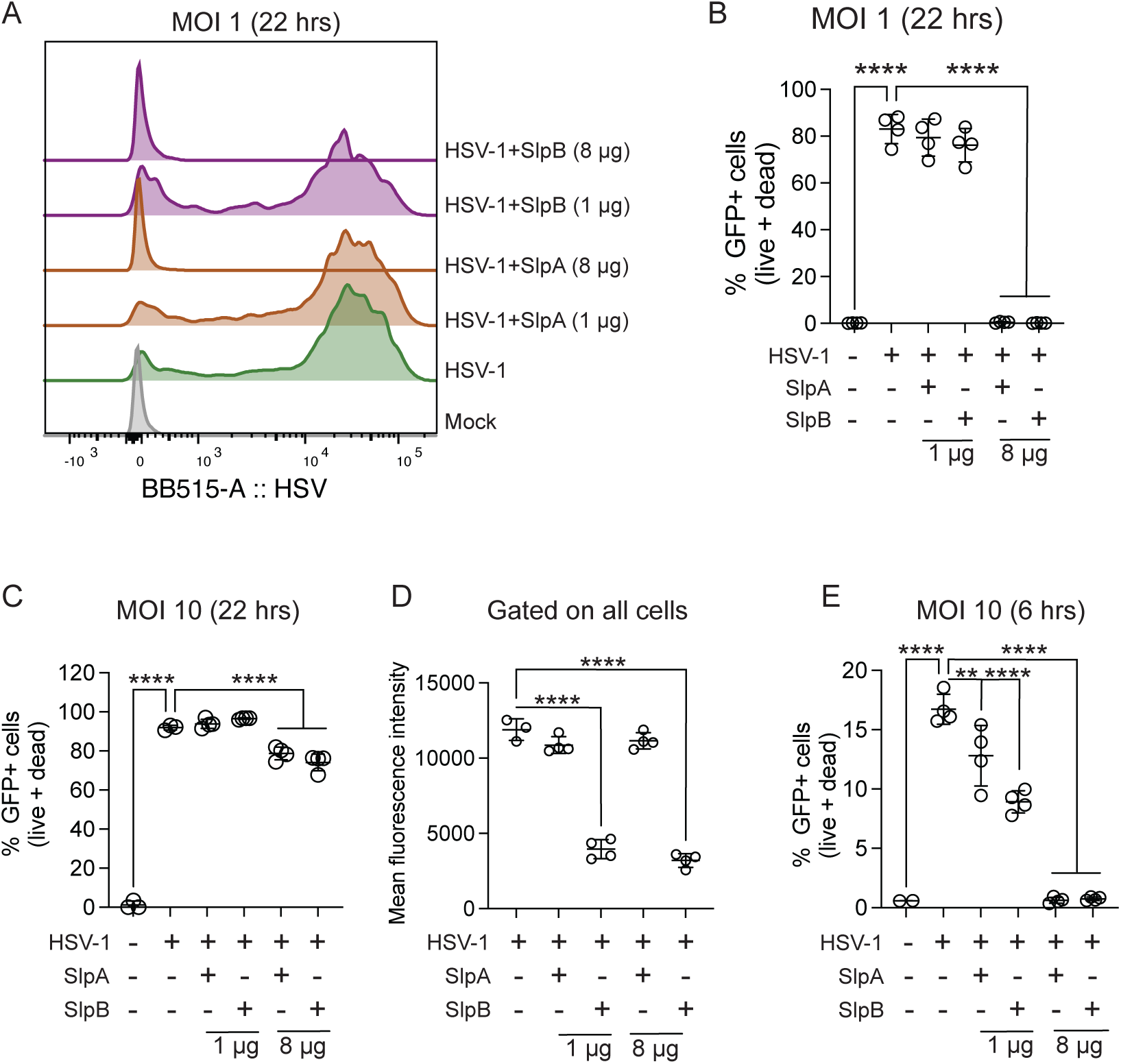
*L. crispatus* SLPs inhibit HSV infections. HFFs were infected with HSV-1 K26GFP (A-E) at an MOI of 1 (B) or MOI of 10 (C-E) alongside indicated concentrations of *L. crispatus* SlpA and B, n=4. GFP fluorescence is shown from representative conditions (A) and GFP+ live and dead cells were quantified as a frequency of total cells (B, C, and E). GFP mean fluorescence intensity from (C) was quantified in (D). Data were compared using a 1-way ANOVA with Dunnett’s correction (B-E). Error bars represent mean and SD.

### *L. crispatus* surface layer proteins bind herpes simplex 1 virions

SLPs have been reported to bind glycans (Tajadura-Ortega et al., 2025), proteins (Decout et al., 2024), and collagen (Muscariello et al., 2020) with diverse effects on bacterial and host cell signaling. Thus, we wanted to test if surface layer proteins could directly bind HSV virions. We incubated *L. crispatus* SlpA or SlpB proteins tagged with AF647 along with HSV-1 K26GFP for four hours and evaluated virions using flow virometry (Fernandes et al., 2025) (Figure 6 and Supplementary Figure 8). Replication-deficient HIV virions expressing iGFP were used as a control virus to test if binding was unique to HSV-1. We found that both SlpA (Figure 6A and C) and SlpB (Figure 6B and D) bound HSV-1 virions in a dose-dependent manner while no binding was observed with HIV virions. At the higher concentration (1000 ng), SlpA bound 12.8% of HSV-1 virions in contrast to 1.5% of HIV virions (Figure 6C). SlpB bound HSV-1 virions at a similar frequency with 17% of the virions fluorescing AF647+, a significantly higher frequency when compared to HIV virions (Figure 6D). Together, with the aforementioned infection inhibition data, these data indicate that *L. crispatus* SLPs directly bind to HSV-1 virions in a virus-specific manner, and that this interaction may drive inhibition of infection.

**Figure 6:**
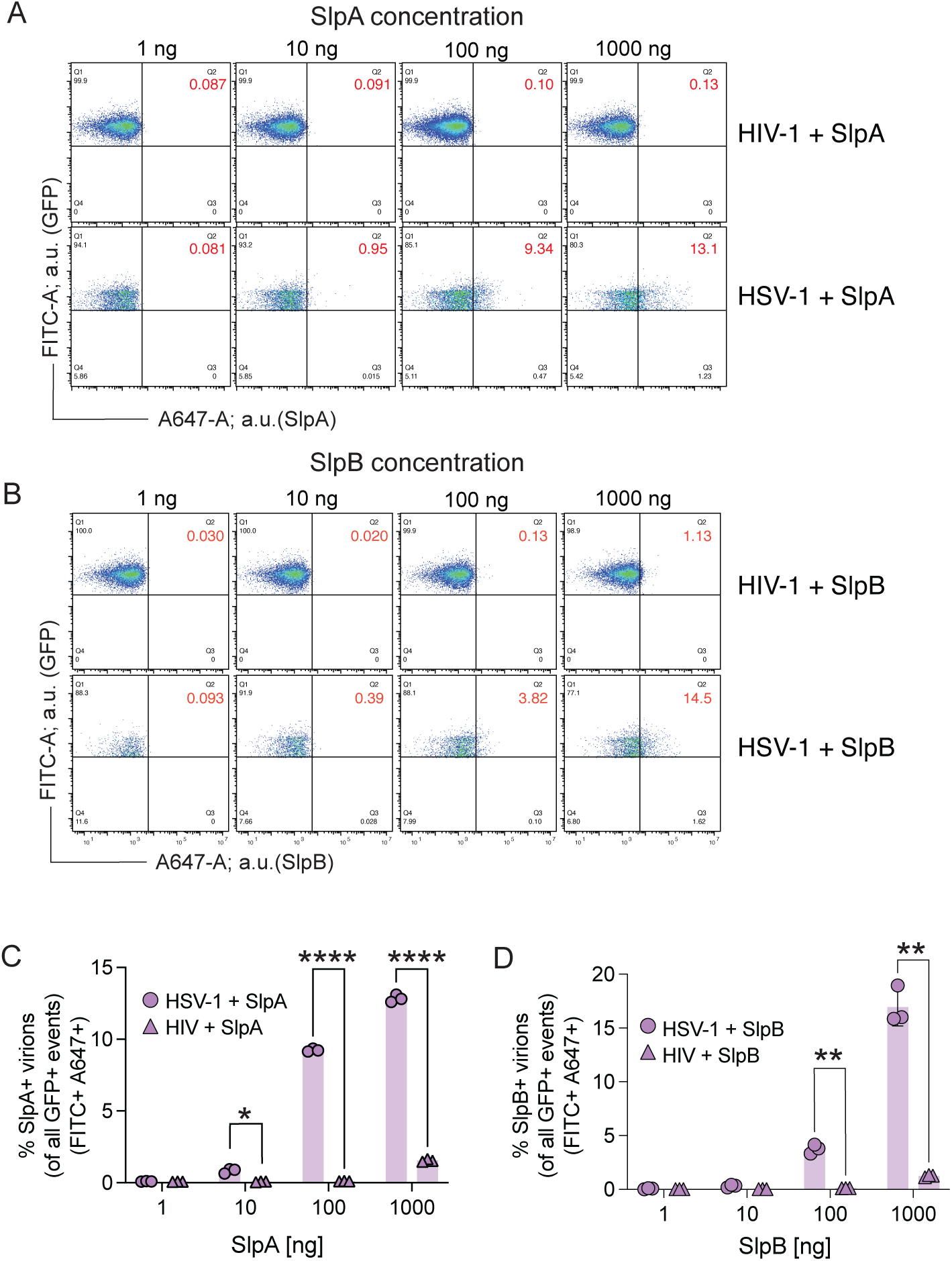
*L. crispatus* surface layer proteins bind herpes simplex virions. (A-D) HSV-1 K26GFP and HIV iGFP virions were incubated with indicated amounts of surface layer proteins SlpA and B for 3.5 hours at 37°C, fixed and run on a flow cytometer optimized for small particle detection. SlpA and B were labeled with AF647 and HSV-1 K26GFP+/AF647+ events quantified. (C-D) AF647+ virions were quantified as a frequency of total virions. Datasets were compared using Welch’s t-test. Error bars represent mean with SD.

### LTA and PG inhibition of HSV infection is independent of TLR2 signaling

Next, we wanted to test the role of cell-intrinsic immune signaling in PG and LTA antiviral protection. Pattern recognition receptor Toll-Like Receptor 2 (TLR2) is known to be activated by lipopeptides including LTA (Travassos et al., 2004; Morath et al., 2002; Henneke et al., 2005). Commercially available PG is also known to activate TLR2 although studies have shown that highly purified PG does not activate TLR2 but rather is sensed by intracellular sensors (Travassos et al., 2004). We first asked if activating TLR2 signaling was sufficient to protect against HSV-1 infection (Figure 7A-B).

**Figure 7:**
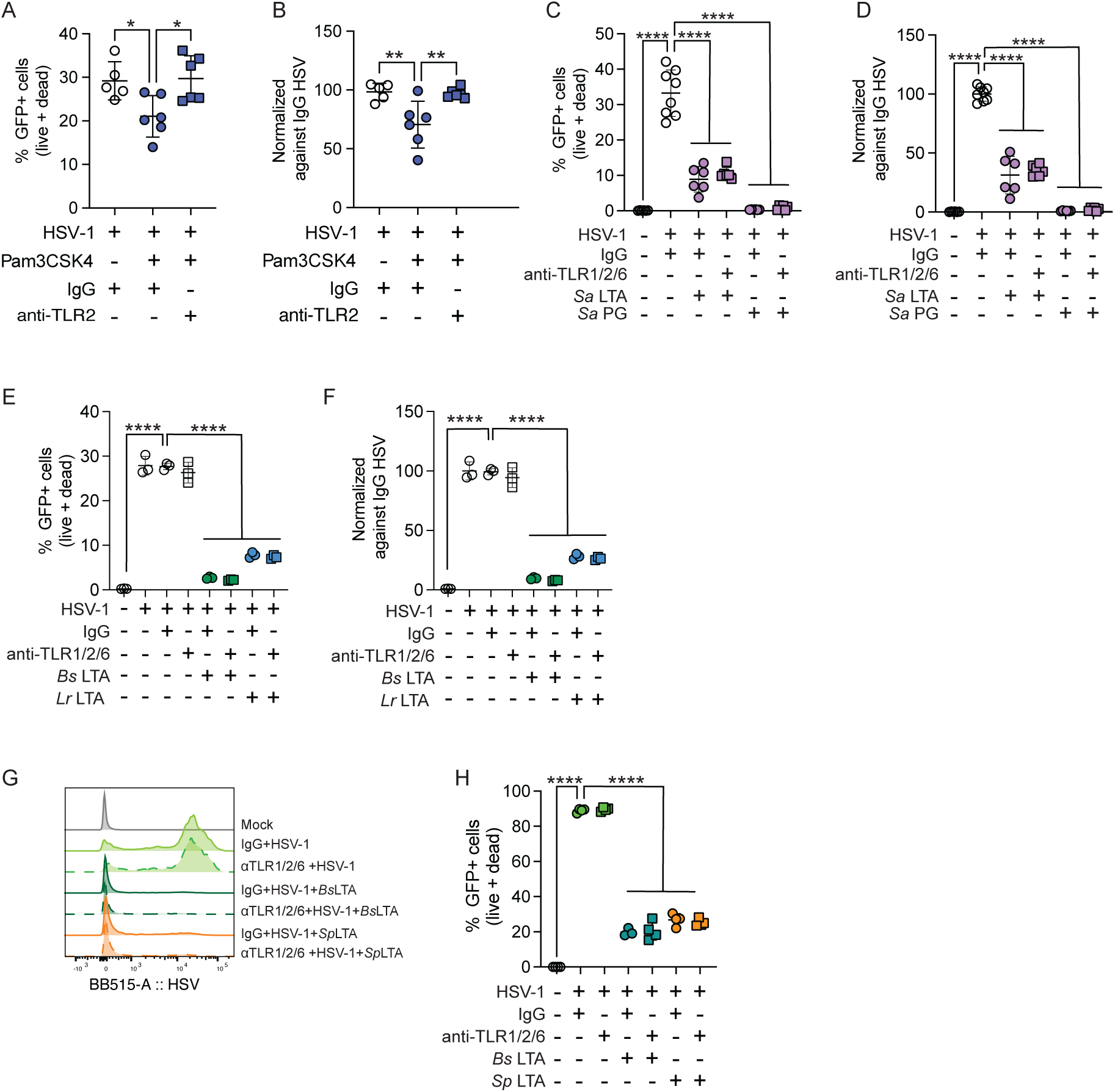
LTA and PG inhibition of HSV infection is independent of TLR2 signaling. (A-E) End1 cells were pre-treated with 0.5 µg each of anti-TLR1, 2, and 6 antibodies or control IgG for 2 hours then infected with HSV-1 K26GFP at an MOI of 1 in the presence of 500 ng Pam3CSK4 (A-B), 50 µg *S. aureus* LTA (InvivoGen) or PG (InvivoGen) (C-D) or 10 µg *B. subtilis* (InvivoGen) or *L. reuteri* LTA (E-F). GFP+ live and dead cells were quantified 22 hours post-infection (A, C, and E) and data normalized against HSV-1 infected cells alone (B, D, and F). (G-H) HFFs were treated with anti-TLR1, 2, and 6 or control IgG antibodies for 2 hours and then infected with HSV-1 K26GFP at an MOI of 1 in the presence of 50 µg LTA from *B. subtilis* (Sigma-Aldrich) or *S. pyogenes* (Sigma-Aldrich), (n=4, a representative experiment is shown). GFP fluorescence profiles from representative samples are shown (E) and quantified (F) 22-hour post-infection. Data were compared using 1-way ANOVA with Tukey’s correction (A-B) or Dunnett’s correction (C-F and H). Error bars represent mean and SD.

Treatment of End1 cells with 500 ng of TLR2-activating synthetic lipopeptide Pam3CSK4 resulted in a small but significant decrease in the frequency of HSV-1 K26GFP infected cells, and this was obviated by pretreatment with TLR2 blocking antibody (Figure 7A-B). Pretreatment with TLR2 blocking antibody did not change the antiviral efficacy of *S. aureus* PG or LTA (Figure 7C-D), indicating that while TLR2 activation could confer minor reductions in HSV infection levels, TLR2 is not required for *S. aureus* PG or LTA antiviral activity in End1 cells. TLR2 forms heterodimers with TLR1 and TLR6 recognizing diacylated and triacylated LTAs (Takeuchi et al., 2002, 2001). Since we do not know the structure of *L. reuteri* LTA, we pretreated End1 cells with antibodies against TLR1,2 and 6 and evaluated the antiviral efficacy of *B. subtilis* and *L. reuteri* LTA (Figure 7E-F). We found that signaling from all three TLRs were dispensable for LTA antiviral activity. We confirmed that signaling via TLR1/2 and TLR2/6 heterodimers are also not required for *B. subtilis* LTA or LTA from another gram-positive vaginal and throat commensal, *S. pyogenes* in HFFs (Figure 7G-H). These data suggest that cell intrinsic immune signaling via relevant TLRs are not required for PG or LTA suppression of HSV-1 infection in barrier cells.

## Discussion

Although bacterial interactions with multiple eukaryotic pathogenic RNA viruses including influenza virus (Rowe et al., 2019), poliovirus (Kuss et al., 2011; Robinson et al., 2014), coxsackie B3 virus (Robinson et al., 2019), reovirus (Kuss et al., 2011; Berger et al., 2017), norovirus (Jones et al., 2014), aichi, and mengo picornaviruses (Aguilera et al., 2019) have been previously described, all of these interactions enhanced, rather than suppressed viral infectivity. Multiple mechanisms including increased receptor binding (poliovirus) (Robinson et al., 2014) and enhanced thermostabilization of virions (reovirus, poliovirus, aichi and mengo picornaviruses) (Aguilera et al., 2019; Waldman et al., 2017; Berger et al., 2017; Robinson et al., 2014) have been described. To the best of our knowledge, these interactions have not been explored in DNA viruses including herpesviruses (Robinson and Pfeiffer, 2014).

Our findings suggest that multiple gram-positive bacterial cell wall components interact with HSV virions to inhibit infection in cells and in mice. In this paper we characterize the role of the major cell wall components PG and LTA as well as *L. crispatus* surface layer proteins on HSV infectivity across multiple human cell lines and in mice. *B. subtilis* PG and LTA have been previously shown to enhance infectivity of reovirus by enhancing virion stability without affecting host cell entry (Berger et al., 2017). In contrast, we found that *B. subtilis* PG and LTA both significantly reduced HSV infection of host cells and the vaginal mucosa of mice. The mechanisms underlying the differences in the outcomes of bacterial-viral interactions between enteric RNA viruses and the vaginal DNA virus HSV remain unclear. One possible contributing factor is that RNA viruses broadly experience increased rates of premature genome loss and reduced virion stability compared to DNA viruses including HSV which express less genome loss and are stable at much higher temperatures (Smith et al., 2009; Bauer et al., 2015; Waldman et al., 2017; Robledo Gonzalez et al., 2023). A second factor may be the differences in relative microbial loads between the gut and vaginal mucosal surfaces, with the vagina containing both a lower diversity of bacterial species and a lower biomass that may result in reduced rates of bacterial-viral interactions and reduced selection pressure for viral pathogen interaction (Mitchell et al., 2015; Pacha-Herrera et al., 2020). Further work is needed to determine how other DNA viruses interact with bacteria in the vagina and across mucosal surfaces. While DNA viruses form the majority of the vaginal virome (Happel et al., 2020; Jakobsen et al., 2019), future work is required to investigate if vaginal RNA viruses including HIV and zika virus, interact with bacterial cell wall components and if these interactions alter infection outcome.

In our experiments, treatment of *B. subtilis* PG with lysozyme reduced PG protection from HSV- 2 infection. This leads us to hypothesize that PG forms a barrier that prevents HSV from effectively entering cells. Additionally, treatment with *E. coli* PG did not protect mice from HSV-2 infection, suggesting that PG protection from infection is unique to gram-positive bacteria.

Gram-negative bacteria have much thinner PG layers than gram-positive bacteria, suggesting that the lack of protection by *E. coli* PG may be due to smaller chains of PG backbone. However, it’s also possible that the peptide linkages and other modifications that are unique to gram-positive PG may be influencing HSV infection (Vollmer, 2008; Vollmer et al., 2008).

Additionally, it remains possible that there is a co-purifying contaminant in *B. subtilis* PG that is lysozyme sensitive and anti-viral. Similarly, although our composition analyses of *L. crispatus* and *L. reuteri* PG suggest that it is pure PG, there is a possibility that there are other co-purifying components that may be impacting HSV infection. This is corroborated by our data suggesting that WTA is present in our *L. crispatus* and *L. reuteri* PG. To our knowledge, the impact of purified WTA on HSV infection has not been explored.

Our data show that between 15-20% of HSV-1 virions interacted with 1000 ng of SlpA and SlpB proteins. While we did observe an increased tendency of these proteins to self-aggregate at higher concentrations which may affect the frequency of bound virions observed, this aggregation did not result in increased binding of HIV-1 virions supporting an HSV-specific interaction. Furthermore, recent literature shows that SLPs may mask other surface structures on vaginal *L. crispatus* (Decout et al., 2024). Thus, it remains possible that *in vivo*, cell wall structures like PG and LTA are masked by SLPs in some lactobacilli, reducing their antiviral effectiveness in the context of the whole bacterial cell. Given that our data demonstrate that purified PG and LTA are very antiviral, this leads us to hypothesize that both the whole bacterial cell and the cell wall components that are shed from the bacterium into the environment (Reith and Mayer, 2011) play a role in restricting HSV infection at the mucosa.

The role that these interactions may play in humans needs to be further investigated as human herpes simplex viruses do not reactivate or go into latency in infected mice. Our data indicate that in our experimental animal and cell models, robust cell wall protection of primary HSV infection requires cell wall protection at the time of infection. However, the ubiquity of genital HSV infections suggests that the presence of lactobacilli is not sufficient to protect against acute infections (Looker et al., 2015b; a). However, amongst seropositive individuals, a wide heterogeneity of recurrence rates is observed with a subset of people subjected to multiple recurrences annually (Agyemang et al., 2018; Tronstein et al., 2011). These recurrences can’t be predictively modeled by local immune responses (Dhankani et al., 2014) but can be correlated to the presence of BV and the absence of lactobacilli (Cherpes et al., 2003, 2005).

Asymptomatic shedding in the absence of a genital ulcer has also been reported in people (Casto et al., 2024; Johnston et al., 2022). Thus, we speculate that the local vaginal microbiome may play a key role in HSV symptom recurrence rates. Based on the data shown here, we propose a model in which individuals with a mucosal microbiome dominated by gram-negative bacteria, like that seen in BV (Figure 8A), are more likely to see a conversion from asymptomatic low-level viral shedding to ulcer formation, inflammation, and increased disease transmission due to lack of gram positive bacterial cell wall components in the mucosal environment. In contrast, increased PG, teichoic acid, and SLP at the vaginal mucosa in a lactobacillus-dominant microbiome (Figure 8B) reduces genital ulcer formation and HSV transmission by interacting with the virions to inhibit further viral infection. Both SlpA and SlpB can be detected in vaginal swabs from people with *L. crispatus*-dominant vaginal communities (Decout et al., 2024). However, like PG and LTAs, quantification of the amount of SLP in the vaginal mucosa in HSV infected people experiencing asymptomatic shedding compared to people with BV and genital lesions is required to determine if these compounds play a physiological role in reducing HSV recurrence.

**Figure 8:**
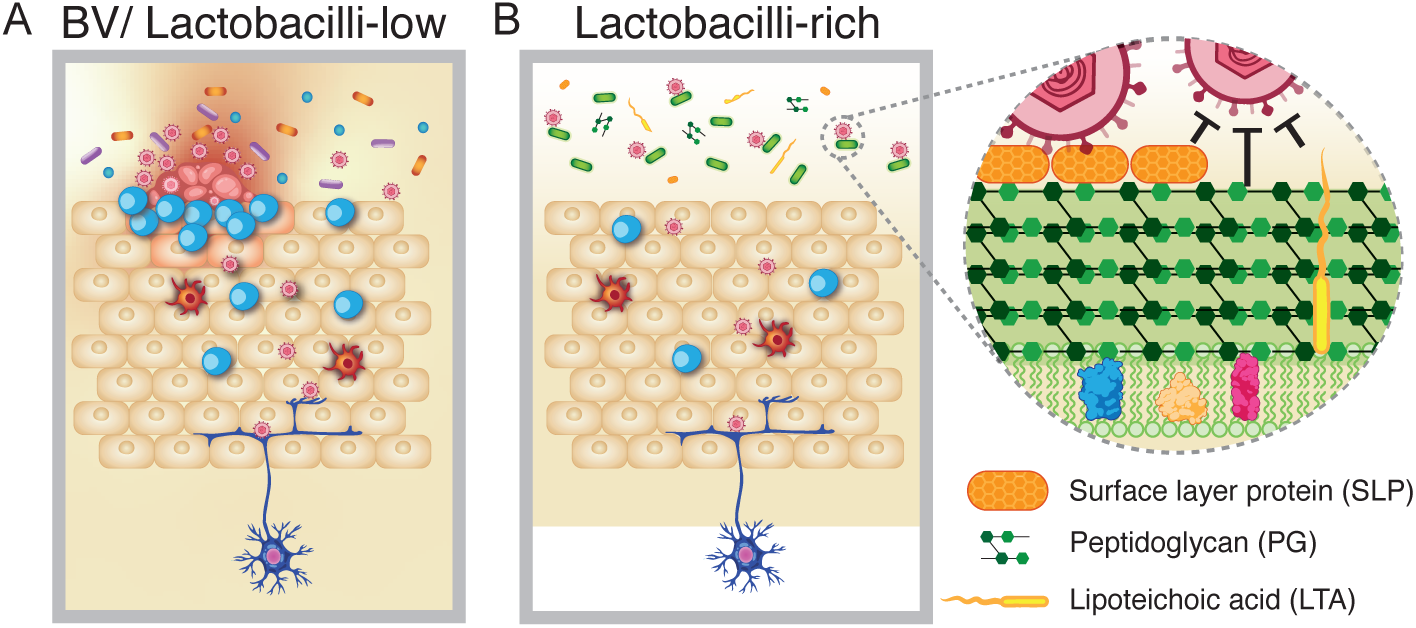
Model of SLP, PG, and LTA inhibition of HSV infection. (A) Gram-negative-rich microbial communities, like that seen in BV, that are low in gram-positive PG, LTA, and SLP increase HSV symptom development risk. (B) Gram-positive-rich microbial communities decrease HSV infection through PG, LTA, and SLP.

The presence of gram-positive bacteria in the skin microbiome, which can include *S. aureus* (used in this study), and the efficacy of PG and LTA in inhibiting HSV infection of foreskin fibroblasts and keratinocytes suggests that gram-positive bacterial cell wall-mediated antiviral activity could be important to the oral and skin mucosal environments as well. Further studies quantifying specific gram-positive cell wall components including free PG, LTA, and SLP in the oral mucosa of asymptomatic seropositive individuals compared to seropositive individuals with multiple recurrences is required to test our hypotheses. Collectively, our data suggest a potential role for gram-positive bacterial cell wall components or killed gram-positive bacteria as a topical application to inhibit herpetic ulcer formation and disease transmission (Gopinath and Adams, 2025).

## Materials and Methods

### Bacterial cultivation

Bacterial strains: *Limosilactobacillus reuteri* CF48-3A (BEI Resources HM-102), *Lactobacillus crispatus* MV-1A-US (BEI Resources HM-637), and *Lactobacillus crispatus* 4M1 and 13M1 (both a kind gift from Julian Marchesi, Imperial College London). Bacteria were cultivated at 37 °C in a standard vinyl anaerobic chamber (Coy Laboratory Products) with input gas of 5% H_2_, 20% CO_2_, N_2_ balance and a gas infuser (Coy Laboratory Products) set to maintain H_2_ concentration between 2.5%-3.0%.

Bacteria were grown in either De Man-Rogosa-Sharpe (MRS) medium (BD Difco cat # 288130 for broth and cat # 288210 for agar) or NYC III medium (5 mM HEPES, 42.8 mM NaCl, 0.5% w/v D-(+)-glucose, 1.5% w/v proteose peptone No. 3, 0.38% w/v yeast extract, 10% v/v heat-inactivated horse serum (Gibco cat # 16050122).

### HSV-2 Vero cell plaque assays

#### General plaque assay protocol

Wild-type HSV-2 186syn+ was used for all HSV-2 infections *in vitro* and Vero cell plaque assays conducted as follows. Six well tissue culture plates were seeded with Vero cells (ATCC CCL-81) and grown to 100% confluency in a monolayer in 1x DMEM + GlutaMAX (+ 4.5 g/L D-glucose + 25 mM HEPES-sodium pyruvate, Gibco cat # 10564-011) supplemented with 10% v/v heat-inactivated fetal bovine serum (HI-FBS) (Gibco cat # 10438-026) and 1% v/v pen/strep (penicillin 10,000 units/ml, streptomycin 10,000 µg/ml, Gibco cat #15140-122). Plaque assays were carried out by serial dilution of HSV-2 mixtures in ABC buffer (PBS supplemented with 1% v/v FBS, 1% w/v glucose, 0.5 mM MgCl_2_ and 0.9 mM CaCl_2_) and infecting the confluent Veros using 250 µl of each dilution per Vero well in the 6 well plate. The viral mixture was pipetted onto the Veros, rocked to disperse, and then allowed to infect the cells under standard tissue culture conditions (5% CO_2_, 37 °C) with rocking every 15 minutes to disperse virus. After one hour of incubation at 37 °C, the ABC buffer was aspirated off the cells and 2 ml/well of 1x DMEM + GlutaMAX supplemented with 1% HI-FBS, 1% pen/strep, and [0.002 mg/ml] human IgG (Sigma-Aldrich cat # I2511) was added. Infections were allowed to continue at 37 °C for 36-42 hours to allow plaques to form before stopping the infection by aspirating the media and staining for plaques with 1% crystal violet in 20% ethanol. Plaques were stained for 30 minutes before stain removal and plaque counting for PFU/ml.

#### Cell wall treatments

For testing the impact of cell wall components on HSV-2 infectivity *in vitro* (Figures 1-3), the following compounds were used: *B. subtilis* PG (Sigma-Aldrich cat # 69554), *S. aureus* PG (Sigma-Aldrich cat # 77140), MurNAc (Sigma-Aldrich cat # A3007), GlcNAc (Beantown Chemical, cat # 13234), lysozyme (from chicken egg white, Sigma-Aldrich cat # L6876), and *L. crispatus* and *L. reuteri* PG (see below for purification details). Commercial cell wall component freezer stocks were diluted to [10 mg/ml] in either pyrogen free water or 1x PBS and stored at -20 °C, whereas *L. crispatus* and *L. reuteri* PG were diluted to [10 mg/ml] in pyrogen free water and stored at -80 °C. Lysozyme was prepared fresh for each assay from a powder stored at -20 °C by dilution into 1x PBS and filter sterilized with a 0.22 µm filter. Cell wall treatments were conducted by mixing the cell wall components and/or lysozyme with 40,000 PFU HSV-2 in 1x PBS to a final experimental volume of 250 µl/well in a 96 well plate. The cell wall component and lysozyme concentrations are listed in the appropriate figures as the final [mg/ml] concentrations in the 250 µl experiment volume. The mixtures were then incubated under standard tissue culture conditions (5% CO_2_, 37 °C) for two hours without additional agitation during incubation. The mixtures were then pipetted to mix and serially diluted -1, -2, and -3 in a final volume of 300 µl in ABC buffer for plating 250 µl of each dilution per well in a 6-well Vero plaque assay using the infection protocol above.

### Bacterial live/dead HSV-2 interaction assays

*Lactobacillus crispatus* MV-1A-US was grown from single colonies at 37 °C in 3 ml of NYC III media in a 15 ml vented snap cap round bottom plastic tube (VWR cat # 60818- 725) (n=3). 24 hours later, cultures were OD-normalized and then sub-cultured 1:1,000 for an additional 24 hours. Cultures were then pelleted at 3,000 *g* for 10 minutes at room temperature, supernatant decanted and pellets suspended in 5 ml PBS and spun down again for a total of 3 washes. Final pellets were suspended in 2 ml PBS and OD-normalized by diluting each strain in PBS to an OD_600_ of 1.0 in 3 ml. For UV killing experiments, 1 ml of normalized bacterial cells was then put into a single well of a 6 well tissue culture treated plate and placed directly underneath a UV bulb in a class II A2 biosafety cabinet for 20 minutes at room temperature with the tissue culture plate lid off. During the UV treatment, the remaining non-UV treated control bacteria were left in an Eppendorf tube in normal room light. After UV treatment, 410 µl of both live and dead bacteria were each mixed with 2 µl of HSV-2 virus [3.61 x 10^8 PFU/ml] in Eppendorf tubes with an HSV-2 only control being 2 µl of HSV-2 virus [3.61 x 10^8 PFU/ml] mixed with 410 µl of PBS. For phenol killing assays, bacteria were normalized to an OD_600_ of 1.0 in 1 ml of PBS as described for UV-killed bacteria. The 1 ml was split in half, with 500 µl sitting at room temperature (live cells) while the other 500 µl was phenol killed by adding 4 µl phenol to 500 µl cells. Samples were incubated for 5 minutes at room temperature and then pipetted to mix to ensure even distribution of phenol. Samples were then incubated further at room temp to a final 10-minute incubation time. After phenol killing, both the control phenol-free tubes and the phenol-killed tubes were washed by adding 500 µl of PBS and pelleting at 4,000 *g* for 5 minutes at room temperature. Supernatant was decanted and pellets washed a second time. Supernatant was then pipetted off and 205 µl of PBS added to suspend the pellets. Then, 205 µl of mix was moved to a new tube and mixed with 1 µl of HSV-2 virus [3.61 x 10^8^ PFU/ml] and flicked to mix. HSV-2 only control was 1 µl of HSV-2 virus [3.61 x 10^8^ PFU/ml] in 205 µl PBS. For both UV-killed assays and phenol-killed assays, the bacteria-HSV-2 mixtures were incubated under standard tissue culture conditions (37 °C, 5% CO2) with mixing by inversion every 15 minutes for one hour. All tubes, including the HSV-2 only control, were then centrifuged at 4,000 *g* for 5 minutes at room temperature. Then, 100 µl of supernatant was pipetted into a 96 well plate, and 30 µl serially diluted in ABC buffer -1, -2, and -3 in a final volume of 300 µl for plating 250 µl of each dilution per well in a 6-well Vero plaque assay. Veros were infected as described above in “general plaque assay protocol.”

### HFF infections and flow cytometry

HFFs (ATCC SCRC-1041) were a kind gift from David Knipe (Harvard Medical school) and cultured in DMEM with 10% FBS. For experiments, 3×10^4^ cells per well were plated, rested overnight, and infected at MOI 0.1, 1, and 10 with HSV-1 K26GFP. Virus and cell wall components were mixed together before addition to cells. To promote infection, virus and bacterial cell wall components were maintained on cells for the duration of the infection. At 6 and 22-24 hours post-infection, cell supernatant and trypsinized cells were pelleted, stained with live/dead stain (Thermo Fisher Scientific, cat # L34973), and fixed (Cytofix, BD cat # 554655) for 20 minutes at 4°C. Cells were washed, brought up in FACS buffer (1% FBS, PBS with 0.1% sodium azide), and run on a FACSymphony A3 (BD Biosciences) flow cytometer. To account for experimental variability, infection frequencies were normalized against HSV-1 infected controls, allowing us to compare antiviral efficacy across experiments. Compounds included *S. aureus* LTA (InvivoGen cat # tlrl-pslta), *S. aureus* PG (Sigma-Aldrich cat # 77140), *S. pyogenes* LTA (Sigma-Aldrich cat # L3140), *B.subtilis* LTA (Sigma-Aldrich cat # L3265), Pam3CSK4 (InvivoGen cat # tlrl-pms), and *B.subtilis* PG (Sigma-Aldrich cat # 69554).

For antibody blocking experiments, 0.5 µg of TLR blocking antibodies or control IgG were added to each well of a 24 well plate for 2 hours before infection and maintained during infection in a total volume of 300 µl per well. Antibodies were obtained from InvivoGen (αTLR1, cat # mabg-htlr1-2; αTLR2, cat # mab2-mtlr2; αTLR6, cat # mabg- htlr6-2 and mouse IgG control, cat # mabg1-ctrlm).

For SlpA/B protein experiments (see protein preparation information below in SLP protein purification and flow virometry), indicated amounts of unlabeled SlpA or SlpB (1 µg [4 µg/ml] and 8 µg [32 µg/ml]) were incubated with HSV-1 at MOI of 1 for 10 minutes at 37°C and then added to HFF cells in 150 µl per well of a 24-well plate. One hour post adsorption, media was added to a volume of 250 µl per well and infection rates were evaluated 22 hours post-infection using flow cytometry as described above.

### *Lactobacillus* PG purification

#### Bacterial pellet preparation

For each strain, *L. crispatus* MV-1A-US and *L. reuteri* CF48-3A, a single colony was grown for 24 hours at 37 °C in 7 ml of MRS media in a snap cap round bottom plastic tube in an anerobic chamber. The culture was suspended and diluted 1:1,000 into 4 x 500 ml polystyrene corning bottles of 500 ml anaerobic MRS media. The 2 L was cultured for 24 hours, pelleted at 4,000 *g* for 15 minutes at 4 °C, re-suspended in equal volumes 1x PBS, and pelleted again at 4,000 *g* for 15 minutes at 4 °C. Pellets were left in 4 °C for 0-2 days before being resuspended in 30 ml of 20 mM sodium phosphate buffer pH 7.2 in a 50 ml conical tube. To kill the bacteria, 600 µl of phenol was added to each conical tube and pipetted up and down to mix. Suspensions were incubated for 5 minutes at room temperature and then pipetted up and down to mix before incubating another 5 min. Suspensions were then pelleted at 4,000 *g* for 15 minutes at room temperature before pipetting off the supernatant. Pellets were then suspended in 30 ml of 20 mM phosphate buffer pH 7.2 to wash. Suspensions were pelleted at 4,000 *g* for 15 minutes at room temperature. Pellets remaining in the 50 ml conical tubes were put into the -80 °C.

#### PG extraction

Following established methods for PG purification (Wu et al., 2013) the cells were suspended in cold PBS buffer and placed in a boiling water bath for 10 minutes. Following this, 4% SDS was added and incubated for 40 min. The suspension was cooled to room temperature and centrifuged at 30,000 rpm and 20 °C for 20 min. Next, the supernatant was removed, the pellet resuspended in water and centrifuged; the pellet washing was repeated 4 times. The final pellet then was resuspended in 90% EtOH, centrifuged, and frozen overnight. Next, enzymatic digestion with benzonase was performed overnight, followed by digestion with proteinase K. The sample was then washed with water 4 times, then 90% EtOH on final wash, and lyophilized. From this, 934 mg of *L. reuteri* and 674 mg *L. crispatus* PG material was obtained and used for Vero and mouse experiments. Additionally, the presence of PG and depletion of LTA was validated using GC-MS and NMR as described below.

### Composition analysis of PG

Compositional analytical protocols were developed as previously described (Parimi et al., 2015).

#### HF treatment

5 mg of the PG samples obtained above were suspended in around 400 ul of 48% HF at 4 °C to remove WTAs, a phosphorylated polymer attached to the PG. The samples in HF were then kept in cold room and stirred gently for 48 hours, then centrifuged at 4 °C and pelleted. The pellets were then washed with water 5 times to remove all traces of HF. The samples were then lyophilized overnight.

#### Mutanolysin digestion

The dry material was dissolved at 4 mg/ml in 50 mM Tris buffer (pH 7.4), mutanolysin was added (500 U), and the suspension agitated at 37 °C overnight. The following day, the same amount of mutanolysin was added and the digestion repeated.

#### Trimethylsilylation (TMS) and GC-MS analysis

Approximately 300 µg of each sample was mixed with 20 µl 1 mg/ml *myo*-inositol as internal standard and lyophilized. The dry sample was suspended in 200 µl 6 M HCl and hydrolyzed for 18 hours at 105 °C. After cooling to room temperature, the sample was dried with a stream of filtered air. Traces of acid were removed by repeated addition and evaporation of water (3 times). The dried sample was mixed with 300 µl 1 M methanolic HCl and heated to 80 °C for 2 hours. The methanol was evaporated with air, and residual acid was removed by repeated addition and evaporation of methanol (3 times). Amino groups were re-N-acetylated by incubating the samples with 400 µl of a 2:1:1 mixture of MeOH, pyridine, and acetic anhydride at 100 °C for 1 hour. The volatiles were evaporated with air, and the samples were treated with 150 µl Tri-Sil (Thermo Fisher Scientific) at 80 °C for 30 minutes. After filtration through glass wool, the samples were concentrated with a stream of air and dissolved in 100 µl hexanes. GC-MS analysis of 1 µl of the TMS-derivatized methyl glycosides was performed on an Agilent 7890A GC interfaced to a 5975C MSD, using an Supelco Equity-1 fused silica capillary column (30 m X 0.25 mm ID). The temperature gradient was 80 °C for 2 minutes, raise temperature to 140 °C at 20 °/minute, hold for 2 minutes, raise temperature to 200 °C at 2 °/minute, raise temperature to 250 °C at 30 °C/minute, and hold at 250 °C for 5 minutes.

#### Heptafluorobutyrate derivatization and GC-MS analysis of amino acids

Hydrolysis and methanolysis were carried out as described under ”Trimethylsilylation and GC-MS analysis” using 2-aminoadipic acid instead of inositol as internal standard. The samples were then HFB-derivatized by addition of 200 µl acetonitrile and 50 µl heptafluorobutyric anhydride (HFBA) reagent (Sigma-Aldrich) and heating to 100 °C for 30 minutes. The derivatized amino acids were dried and extracted in 200 µl fresh acetonitrile. GC-MS analysis of 1 µl of the HFB-derivatized amino acids was performed on an Agilent 7890A GC interfaced to a 5975C MSD, using an Agilent DB-5 fused silica capillary column (30 m X 0.25 mm ID), using the same temperature gradient as described under ”Trimethylsilylation and GC-MS analysis”.

#### NMR Spectroscopy

∼ 1 mg of HF-treated and mutanolysin-treated PG sample was suspended in 500 µl D_2_O (99.9 % D) and lyophilized overnight. The samples were dissolved in 500 µl of DMSO-d_6_ and 200 µl of D_2_O (99.96 % D). The sample was transferred to a 5-mm NMR tube for NMR analysis. NMR data were acquired at 298 K on a Bruker Avance III spectrometer (^1^H, 600 MHz) equipped with a cryoprobe using standard pulse sequences. The acquisition parameters were 64k complex data points, 20 ppm spectral width, 4.7 s total recycle delay and 16 scans. Chemical shifts were referenced to DMSO-d_6_ (δ_H_ = 2.50 ppm). The spectra were processed and analyzed with *MestReNova* v14.2.1-27684.

### Genital HSV infections

All animal experiments were performed under an IACUC-approved protocol ISO0003145 in the Harvard. T. H. Chan School of Public Healthanimal facilities accredited by the Association for Assessment and Accreditation of Laboratory Animal Care (AAALAC) International. Mouse infections were carried out as previously described in (Gopinath et al., 2018; Glick et al., 2024). C57BL/6 mice between 6-8 weeks old (Charles River Laboratories, MA, USA) were subcutaneously injected with 2 mg DMPA (Prasco Laboratories) 5 days prior to viral infection to synchronize estrous cycles to enhance sensitivity to HSV infection (Linehan et al., 2004). Mice were swabbed intravaginally with a PBS-soaked calcium alginate swab (Calgiswab, Puritan) before intravaginal infection with 5,000-10,000 PFU wild-type HSV-2 186syn+ alone or with indicated treatments (see below) in a total volume not exceeding 20 µl per mouse. Vaginal lavage was collected on indicated days by swabbing with a PBS-soaked Calgiswab, pipetting 50 µl of PBS into the vagina, pipetting up and down four times and adding the exudate to 950 µl of ABC buffer. Vaginal viral titers were quantified via plaque assay on Vero cells as described above in “general plaque assay protocol.” Disease progression was tracked for 12-14 days and survival up to 21 days. Disease was scored as follows: 1) Vaginal erythema and inflammation without hair loss present, 2) perianal hair loss present, 3) signs of morbidity presenting including hunched posture, ruffled fur due to a reduction in grooming, 4) hind-limb paralysis, 5) death by disease or by euthanasia. Animals were euthanized if they were unable to access food due to hind-limb paralysis, had lost 20% of their weight from the onset of the experiment, or if they showed excessive morbidity.

#### Bacterial cell-wall vaginal treatments

For all cell-wall treatments, unless noted otherwise, virus was mixed with the indicated cell wall component, enzyme, pyrogen-free water, or PBS right before infection with appropriate vehicle controls used (either water or PBS). All *L. crispatus* and *L. reuteri* PG and LTA treatments were suspended in water. For *L. crispatus* and *L. reuteri* PG treatments at time of infection (Figure 1), mice received 200 µg of either *L. crispatus* or *L. reuteri* PG mixed with 10,000 PFU HSV-2.

Control mice received HSV-2 mixed with PBS (first independent experiment, data shown) or water (repeat independent experiment, data not shown). For *L. crispatus* and *L. reuteri* PG treated mice, mice were re-swabbed and re-treated a second time 24 hours later with 200 µg of PG and control mice treated with equivalent volumes of PBS or water. For mouse experiments in Figure 2, mice received 50 µg of the indicated PG suspended in water or PBS mixed with 10,000 PFU HSV-2, and controls included HSV- 2 mixed with equivalent volumes of appropriate vehicle controls (either water or PBS.) Treatment sources include *B. subtilis* PG (Sigma-Aldrich, cat # 69554), *S. aureus* PG (InvivoGen, cat # tlrl-pgns2), and *E. coli* PG (InvivoGen, cat # tlrl-kipgn). For mouse experiments in Figure 3, mice were infected with 10,000 PFU HSV-2 diluted in PBS with or without 50 µg of *B. subtilis* PG (Sigma-Aldrich cat # 69554) and 10 mM lysozyme (from chicken egg white, Sigma-Aldrich cat # L6876) diluted in PBS. For mouse experiments in Figure 4, mice were infected with 10,000 PFU HSV-2 diluted in water with or without 50 µg of *L. reuteri* LTA diluted in water. For mouse experiments in Supplementary Figure 4, mice were infected with 5,000 PFU HSV-2 and 4 hours post-infection, swabbed and treated intravaginally with 200 µg *L. crispatus* PG or water in a 20 µl final volume. For 5 days post-infection, mice received an additional 200 µg *L. crispatus* PG/20 µl or water daily. On days 6 and 7, mice were treated with a smaller volume due to virus-driven vaginal inflammation closing the vaginal canal. On days 6 and 7, mice received either 100 µg *L. crispatus* PG or water in 10 µl volume.

### Germ-free mouse animal housing and maintenance

10-week-old C57BL/6 germ-free female mice were housed with autoclaved food, water, and autoclaved bedding, which was changed on a weekly basis in a Biosafety Station under germ-free conditions. Germ free conditions were maintained using a Tecniplast isoCage System (Type: ISO36PFEUS) and isoCage Biosafety Station (Type: 9ISODTU4) which were used to house and work with germ-free mice, respectively.

Experiment materials and sealed isoCages were dunked in 3 µg/ml MB-10 sterilant for at least 30 seconds before transferring to the prepared biosafety station. Fecal pellets were collected from each cage every time the isocages were opened in the biosafety cabinet. DNA was extracted from stool samples using the QIAamp Fast DNA Stool Mini Kit (cat. no 51604). The presence of bacterial genetic material, or lack thereof, was confirmed via qPCR using 16S primers. A cycle threshold value of 28 and above was used to confirm germ-free status.

### Germ-Free mouse infections

Germ-free mice, kindly provided by Wendy Garrett (Harvard T.H. Chan School of Public Health), were injected subcutaneously with 2 mg DMPA to synchronize estrus cycles and increase susceptibility to viral infection. Seven days post-injection, mice were infected intravaginally with 10 µl containing 5,000 PFU HSV-2 and either 100 µg of *L. crispatus* PG or additional volume of PBS. The stock solution of *L. crispatus* PG used in this infection was [10 mg/ml] suspended in water. To reduce contamination risk, vaginal lavage was collected only on day 2 post-infection. Vaginal viral titers were quantified using plaque assays on Vero cells. Disease progression was tracked daily over the course of 14 days and survival was tracked for 21 days as described above. Disease scores were recorded starting on day 6 post-infection. Scores were not collected on day 10 or 12 post-infection to reduce the risk of contamination.

### Primary human keratinocyte culture and infections

NHEK were isolated from discarded human foreskin tissue in accordance with the University of Massachusetts Chan Medical School Institutional Review Board #H00021295. Keratinocytes were isolated from the tissue as previously described (Orzalli et al., 2021) and initially cultured on mitomycin-treated 3T3. Prior to experimentation, NHEK were transferred to Keratinocyte Serum Free Media (KSFM) supplemented with epidermal growth factor (EGF) and bovine pituitary extract (BPE) (ThermoFisher).

#### Virus Infections

NHEK were plated in KSFM at 3 x 10^4^ cells/well of a 96 well plate the day prior to infection. HSV-1 K26GFP (Desai and Person, 1998) was propagated and viral titers determined in Vero cells. For NHEK infections, virus was diluted in KSFM with or without *S. aureus* PG (1, 0.25mg/ml diluted in sterile water) (InvivoGen cat # tlrl-pgns2) in a round bottom 96 well plate and incubated at 37°C for 2 hours, after which time it was incubated with the cells for 1 hour at 37°C with gentle shaking. Following the adsorption period, virus containing media was removed and replaced with fresh KSFM containing Hoechst 33342 (Invitrogen) diluted to a final concentration of 3 µg/ml. Cells were imaged hourly for 24 hours using a Lionheart FX automated microscope (Biotek). Image analysis was performed using Gen5 software and ratios of GFP+ to Hoechst+ cells were plotted as a measurement of the percentage of HSV-1 K26GFP infected cells. The area under the curve was determined for each condition and statistical analysis was performed in PRISM.

### LTA isolation and detection

#### Extraction from cells

*L. reuteri* CF48-3A and *L. crispatus* MV-1A-US bacterial pellets were generated as done for PG isolation (see above) and LTA isolation carried out (Han et al., 2003). Pellets were then lyophilized and weighed. 1.46 g of *L. reuteri* cells and 1.32 g of *L. crispatus* were then suspended in 10 ml of 100 mM Na citrate, pH 4.7, followed by addition of 10 ml 1-butanol, and the mixture was shaken vigorously (280 rpm) for 30 minutes at room temperature. The suspension was centrifuged for 20 minutes at 13,000 g, and the bottom yellow water-rich phase was removed and centrifuged again (20 minutes at 13,000 g). The yellow phase was transferred to another bottle and centrifuged again (20 minutes at 13,000 g). The yellow phase (∼340 ml) was dialyzed in a 1 kDa molecular weight cut-off bag against 2 x 4 L 20 mM Na citrate, pH 4.7 overnight at 4 °C. Finally, the retentate was lyophilized.

#### Hydrophobic interaction chromatography

Three buffers were used: Buffer L (25 mM Na acetate, pH 4.5, 15 vol% 1-propanol), Buffer A (100 mM Na citrate, pH 4.7, 15 vol% 1- propanol) and Buffer B (41 mM Na citrate, pH 4.7, 65 vol% 1-propanol). Octyl-sepharose resin (25 ml) in a 16x200 mm gravity column was equilibrated with Buffer A. All of the dry extract was dissolved in 5 ml of Buffer L and centrifuged at 4,200 *g* for 20 minutes to pellet insoluble material. The sample was loaded on the column by gravity at ∼0.3 ml/minutes, the resin was washed with 100 ml Buffer A (∼0.5 ml/minute) and connected to an HPLC system (Agilent 1260 Infinity II). The material was eluted at 0.2 ml/minute with 200 ml linear gradient from 100% Buffer A to 100% Buffer B, and 5 ml fractions were collected. Fractions containing LTA, as determined by phosphorus assay, were pooled and dialyzed against 1% acetic acid (3 x 4 L) and then water (3 x 4 L), and the retentate was lyophilized.

#### Total phosphorus assay

Selected C_8_-sepharose fractions, together with 0, 30, 60, 100, 150 and 200 nmol KH_2_PO_4_ standards, were analyzed for total phosphorus content. An aliquot of 50 μl of each fraction or standard was mixed with 200 μl 10% H_2_SO_4_ in a 5 ml glass tube and heated at 200 °C for 1 hour. After cooling, 50 μl 30 % H_2_O_2_ was added and the content heated at 200 °C for 40 minutes. To each cooled sample, 980 μl reagent consisting of 0.25% (w/v) Na_2_MoO_4_.2H_2_O and 0.5% (w/v) sodium ascorbate was added and the solution incubated at 45 °C for 20 minutes. Absorbance at 490 nm was determined for each sample using a microplate reader.

#### NMR spectroscopy to determine LTA purity

Several mg of the dry material was dissolved in 500 μl D_2_O (99.9% D), lyophilized, dissolved in 500 μl D_2_O, and transferred into a 5 mm NMR tube. NMR data were collected at 25 °C on a Bruker Avance III (1H, 600.13 MHz) equipped with a cryoprobe. 1H data were acquired with a spectral width of 12 ppm, 65k complex data points, and 16 transients. Prior to the Fourier transformation, NMR data were apodized with an exponentially decaying function (lb = 0.3 Hz), and the baseline of the spectra was corrected automatically using a 3rd-order polynomial. The ^1^H was referenced to the respective water signals at 4.7 ppm. Based on these NMR and total phosphorus results, fractions 17-23 (see Supplementary Figure 7) of *L. reuteri* LTA were used for subsequent cell culture and mouse experiments.

### Endocervical cell culture and infections

End1/E6E7 cells (ATCC CRL-2315) were a kind gift from Caroline Mitchell and cultured in Keratinocyte Serum Free media supplemented with [0.1 ng/ml] recombinant epidermal growth factor and [0.05 mg/ml] bovine pituitary extract. For experiments, 2 × 10^5^ cells per well were plated in a 12-well plate, rested overnight and infected at an MOI of 1 with HSV-1 K26GFP. 50 µg of indicated LTA and PG were incubated with HSV-1 for 37°C for 10 minutes before adding to End1 cells at a volume of 250 <u>µl per</u> <u>well</u>. One hour after adsorption, media was added to a volume of 350 µl. To promote infection, virus and bacterial cell wall components were maintained on cells for duration of infection Between 20-22 hours post-infection, cell supernatant and trypsinized cells were pelleted, stained with live/dead stain, fixed and run on the flow cytometer as previously described above. To account for experimental variability, infection frequencies were also normalized against HSV-1 infected controls allowing us to compare antiviral efficacy across experiments. Antibody blocking experiments were conducted as previously described in HFF infections.

### S-layer protein purification

Extraction of S-layer proteins was performed as previously described (Decout *et al*., 2024). Briefly, *L. crispatus* 4M1 and 13M1 bacterial pellets of 50 ml overnight cultures were washed with PBS and resuspended in LiCl 5M at 4 °C for 15 minutes under stirring. The supernatants were harvested by centrifugation for 10 minutes at 3,000 x *g* and dialyzed against water overnight at 4 °C in Snakeskin dialysis tubing cut off 10 kDa (68035, Thermo Fisher Scientific). Precipitated S-layer proteins were recovered by centrifugation at 20,000 *x g* for 20 minutes. The proteins were solubilized in LiCl 5M and further purified by size exclusion chromatography using Sephacryl S200R (S200HR-250ML, Sigma-Aldrich). The S-layer proteins were run in SDS-PAGE and stained with coomassie blue as shown in Decout et al, 2024 (see Figure 4A.) A single band for SlpA (45 kDa band) was obtained from 13M1 and a single band of SlpB obtained from 4M1 (60 kDa band).

### SLP labelling

The purified SLPs (1 mg/ml) were incubated with 10 equivalents of Alexa Fluor 647 NHS Ester (Succinimidyl Ester) (A20006, Thermo Fisher Scientific) in Lithium Chloride 5M for 1 hour at room temperature. Unreacted Alexa Fluor 647 NHS Ester was removed by ultrafiltration using Vivaspin sample concentrators 10 kDa (Thermo Fisher Scientific cat # 88517).

### SLP-virus flow virometry

Prior to experimental use, Alexa Fluor 647-conjugated SLP samples were stored in LiCl at room temperature. We performed buffer exchange to PBS using an Amicon Ultra Centrifugal Filter, 3 kDA MWCO (MilliporeSigma, cat # UFC500324). Prior to buffer exchange, the filter was pre-wet with 500 µl PBS and spun down at 14,000 *g* for 20 minutes. 50 µl of protein was then added to the filter unit and spun down at 14,000 *g* for 30 minutes. The filter was then inverted and spun down at 2,000 *g* for 2 minutes. The flow-through was then stored at 4 °C. Buffer exchanged proteins were used for experiments within 2 days. All sample preparations were done in 1.5 ml microcentrifuge tubes (Westnet, cat # MCT-150-C-S). SLP was first diluted to 500 ng/µl, 50 ng/µl, 5 ng/µl, and 0.5 ng/µl with PBS. 2 µl of each dilution was then added to 16 µl of PBS in triplicate. To each of these tubes, either 2 µl of 6.5 x 10^4^ PFU/µl HSV-1 K26GFP were added for a final concentration of 1.3 x 10^5^ PFU per sample or 2 µl of ∼10^7^ particles/ml of HIV-1 iGFP Env-deficient molecular clone (BEI Resources, cat # HRP-12455) were added as a labeling control. Samples were then incubated at 37 °C for 3.5 hours.

Samples were then fixed with a 4% paraformaldehyde solution (Beantown Chemical, cat # 140770-10X10ML, lot # 50077008) and put on ice for 20 minutes before additional dilution with PBS for flow virometry. Flow virometry was performed using a Beckman Coulter CytoFLEX S with a standard optical configuration. Samples were acquired for 1- 2 minutes at a sample flow rate of 10 μl/min using the tube loader. Experiments were conducted with a threshold of 1,250 on the violet SSC-H, using the following gain settings: violet SSC, 200; FITC, 2,500; and APC, 3,000.

### Statistical analysis

Datasets were evaluated using GraphPad Prism version 10.6.1, GraphPad Software Boston, MA, USA. P-values: **** (<0.0001), *** (<0.001), **(<0.01), *(<0.05). All *in vitro* experiments were repeated a minimum of 2 times, with a minimum of 3 technical replicates except for a subset of SLP inhibition experiments in Figure 5C-E which were conducted once. All animal infection experiments were conducted a minimum of 2 times with a minimum of 4 animals per condition except for experiments shown in Supplementary Figure 3 and Supplementary Figure 4 which were experiments that were conducted once but had n=7-10 animals per condition.

## Funding, acknowledgements, and disclosures

The authors thank Wendy Garrett and members of the Garrett lab (HSPH) for helpful feedback, resources and breeding germ-free animals for infection experiments. We thank Byron Roman and the members of the labs of Flaminia Catteruccia, (HSPH) for helpful feedback and resources. A.N.D.A. was supported with travel awards to present work in this publication by the HSPH Postdoctoral Association and the Infectious Diseases Society for Obstetrics and Gynecology. J.B. was supported by a REDI fellowship funded by the Canadian Institutes of Health Research. The Complex Carbohydrate Research Center, University of Georgia, was supported by the U.S. Department of Energy, Office of Science, Basic Energy Sciences, Chemical Sciences, Geosciences and Biosciences Division, award # DE-SC0015662 and by the National Institutes of Health (NIH) grant # R24GM137782. J.P. was supported by NIH training grant # T32AI007349 and NIH award # F30AI18891. M.H.O. was supported by NIH award # R01AI182052 and as an Investigator in the Pathogenesis of Infectious Disease (Burroughs Wellcome Fund). K.S.C. was supported by Chan Zuckerberg Initiative Science Diversity Leadership grant # 2022-310965, Howard Hughes Medical Institute Freeman Hrabowski Scholars grant, start-up funds from HSPH, and in-kind gifts of lab equipment, consumables, and supplies from Corning, Inc. S.G. was supported by start-up funds from HSPH, Searle Scholars Program, Blavatnik Biomedical Accelerator at Harvard University, and NIH award # R21AI180508. The authors disclose that the research described in this article is related to a U.S. Patent Application (Serial No. US18/890,271; Publication No. US20250090628A1).

## CRediT author statement

According to the CRediT taxonomy, the authors made the following contributions: A.N.D.A. contributed to conceptualization, data curation, formal analysis, investigation, methodology, project administration, supervision, validation, visualization, writing– original draft, and writing–review and editing. L.E.G. and J.B. contributed to data curation, formal analysis, investigation, methodology, project administration, validation, visualization, writing–original draft, and writing–review and editing. J.P. contributed to data curation, formal analysis, investigation, validation, visualization, and writing–review and editing. A.C. contributed to formal analysis, investigation, methodology, and validation. V.J.G., M.G., and D.R.B. contributed to formal analysis, investigation, and methodology. C.H.K. contributed to formal analysis, investigation, and writing–review and editing. M.A. contributed to investigation, validation, and writing–review and editing. M.J. contributed to investigation and validation. G.K. and J.L.C. contributed to formal analysis and investigation. A.K. and L.D.F. contributed to formal analysis, investigation, and validation. J.V. contributed to formal analysis, methodology, supervision, visualization, writing–original draft, and writing–review and editing. M.H.O. contributed to data curation, formal analysis, project administration, resources, supervision, and writing–review and editing. K.S.C.-H. and A.D. contributed to resources, supervision, and writing–review and editing. P.A. contributed to conceptualization, funding acquisition, project administration, resources, and supervision. S.G. contributed to conceptualization, data curation, formal analysis, funding acquisition, investigation, methodology, project administration, resources, supervision, validation, visualization, writing–original draft, and writing–review and editing.

---

**Supplementary Figure 1:**
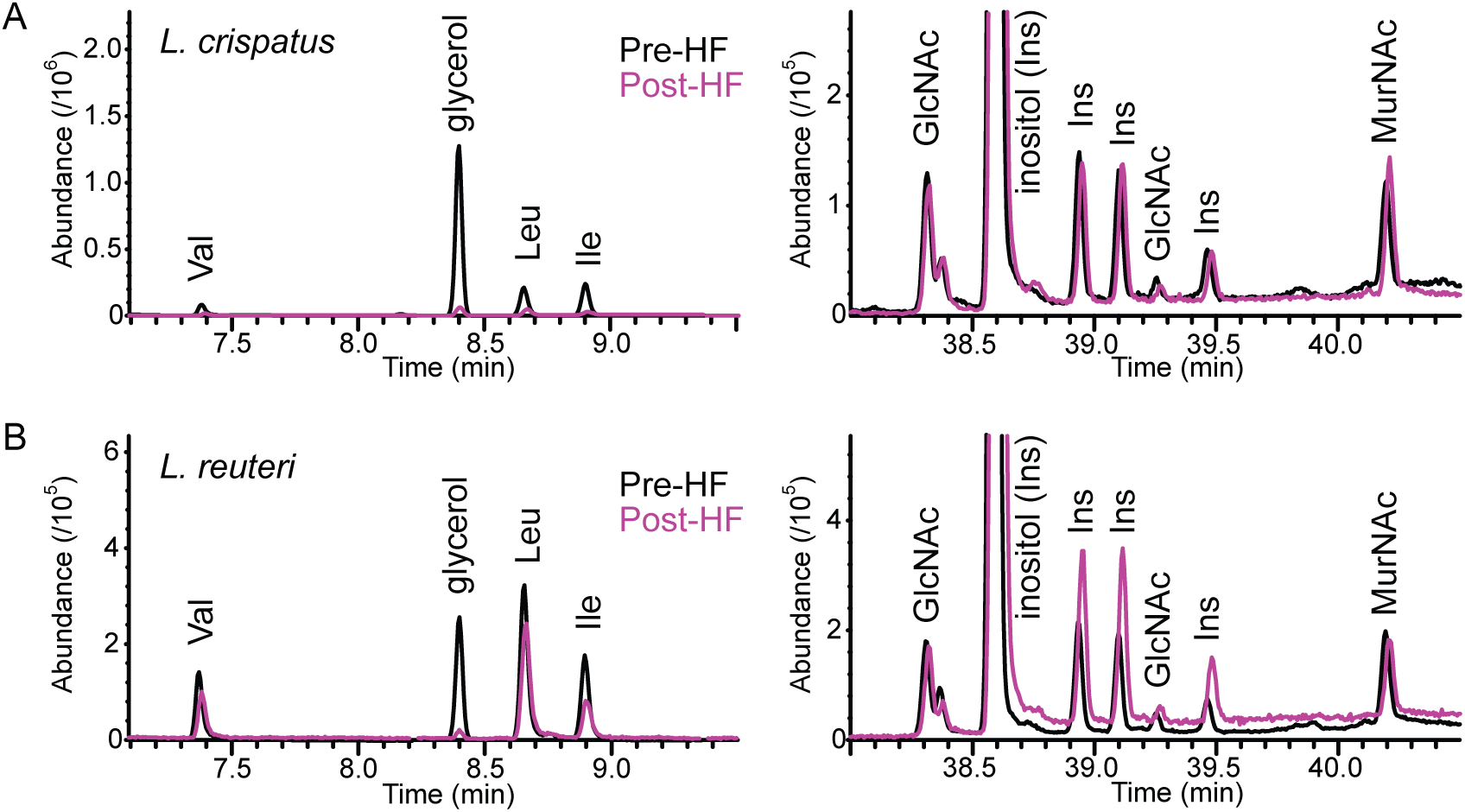
Partial GC-MS chromatograms of TMS derivatives of *L. crispatus* and *L. reuteri* PG before and after HF treatment. The peak of glycerol (8.4 min) was greatly reduced upon HF treatment in both (A) *L. crispatus* and (B) *L. reuteri*, suggesting removal of glycerol phosphate polymer, such as a WTA. Besides glycerol, this region contained peaks due to TMS derivatives of amino acids as indicated. The intensities of the two chromatograms for each bacterial PG were normalized to give approximately the same intensities of GlcNAc- and MurNAc-derived peaks in the pre- and post-HF chromatograms, as shown in the region 38–41.5 min on the right.

**Supplementary Figure 2:**
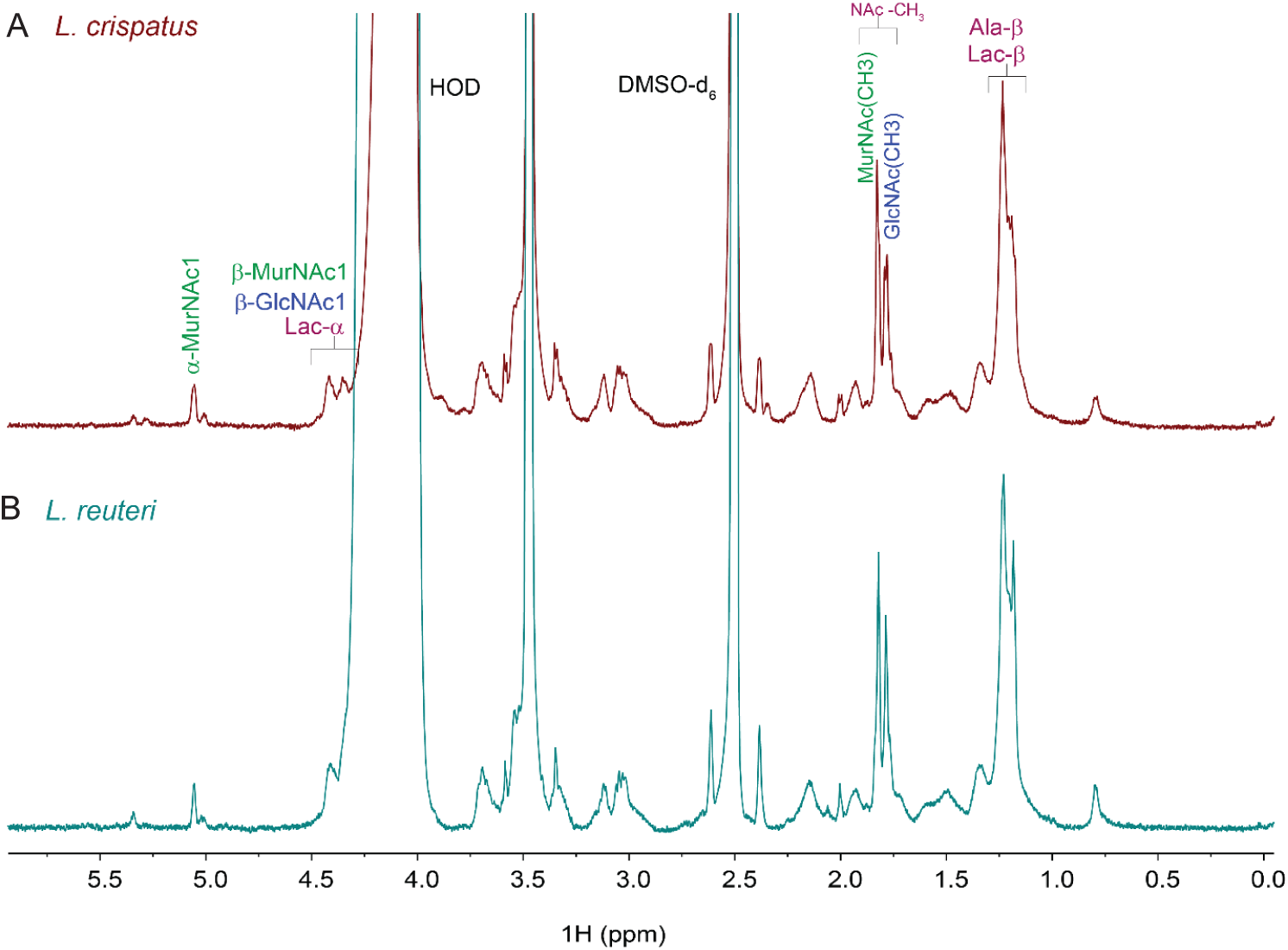
1D Proton NMR spectra of HF-treated and mutanolysin digested *L. crispatus* and *L. reuteri* PG. 1D ^1^H NMR spectrum of **a**) *L. crispatus* and **b**) *L. reuteri* PG (in 3:1 DMSO-d_6_/D_2_O). Peak identification is based on the previously reported ^1^H NMR chemical shifts for bacterial PG in DMSO-d_6_ (Klaić, et al 1990, Fehér K et al. 2003). The peaks at 4.41-4.35 ppm are consistent with the β-anomeric forms of GlcNAc and MurNAc, as well as lactyl H-α group of MurNAc (Lac-α). Both *L. crispatus* and *L. reuteri* PG showed distinct ^1^H signal at 5.05 ppm, consistent with H1 of the reducing-end α-MurNAc, which is a product of mutanolysin digestion. N-acetylation is evident by the peaks at 1.7-1.8 ppm which are due to the methyl protons of N-acetyl groups from β-GlcNAc and β/α-MurNAc residues. The intense peak at 1.2 ppm was tentatively assigned to overlapping signals from lactyl-β and alanine-β methyl groups. Other peaks between 0.5 and 4 ppm are likely due to amino acids. These observations in ^1^H NMR of the isolated material from *L. reuteri* and *L. reuteri* support the presence of the partially digested PG.

**Supplementary Figure 3:**
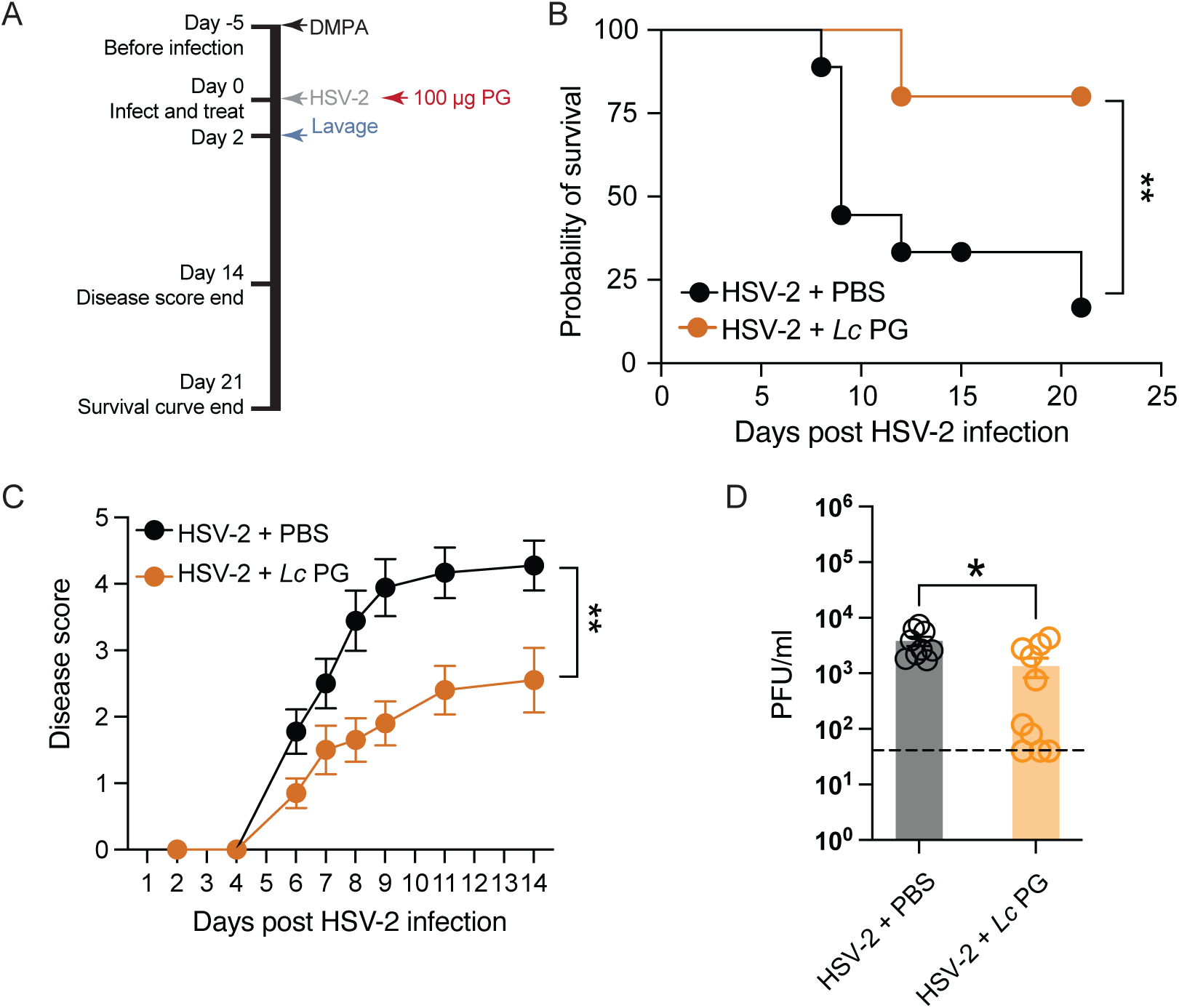
*L. crispatus* PG suppresses HSV-2 infection independently of the host microbiome. (A) Schematic of mouse treatment timeline for the experiments shown in (B-D). Germ-free C57BL/6 mice were injected subcutaneously with 2 mg DMPA to synchronize estrous cycles (n=9 for PBS and n=10 for *Lc* PG. Five days later, they were infected intravaginally with 5,000 PFU HSV-2 and either 100 µg of *Lc* PG or equivalent volumes of PBS (HSV-2 only) at the time of infection. Survival was tracked for 21 days (B) and disease progression was tracked for 14 days (C). (D) Vaginal lavage was taken on day 2 post-infection and viral titers were quantified by plaque assays on Vero cells. (B-C) Datasets were compared using log-rank (Mantel-Cox) test (B), two-way ANOVA with Geisser-Greenhouse correction (C), and fisher’s exact unpaired t-test (D). Error bars represent the mean with SEM.

**Supplementary Figure 4:**
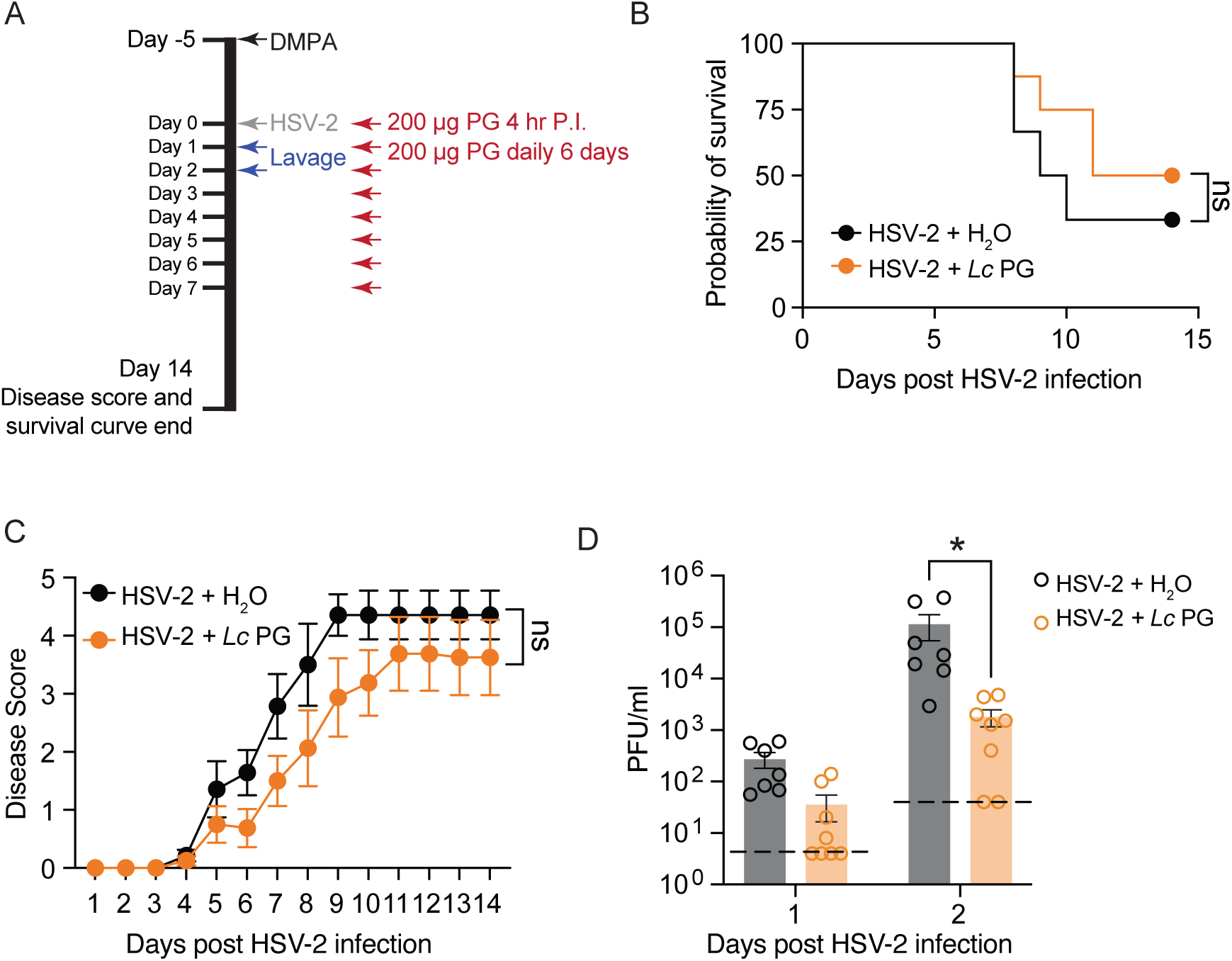
Robust *L. crispatus* PG inhibition of HSV-2 requires PG at time of infection. (A-C) C57Bl6 mice were injected with DMPA to synchronize estrous cycles (n=7 for H_2_O and n=8 for *Lc* PG) and then five days later were infected intravaginally with 5,000 PFU HSV-2 and 200 µg *Lc* PG 4 hours after infection and daily for days 1-5 post-infection. On days 6-7 post-infection, 100 µg of *Lc* PG was used for treatment due to inflammation narrowing the vaginal canal and amount of treatment volume that could be inserted. Survival was tracked over 15 days (B), and disease severity tracked for 14 days (C). (D) Vaginal lavage was taken for 2 days after the time of infection (starting 24 hours after infection) and lavage viral titers quantified using plaque assays on Vero cells. (B and D) Datasets were compared using log-rank (Mantel-Cox) test (B) and two-way ANOVA (C-D) with Geisser-Greenhouse’s correction (C) or Sidak’s correction (D). Error bars represent the mean with SEM.

**Supplementary Figure 5:**
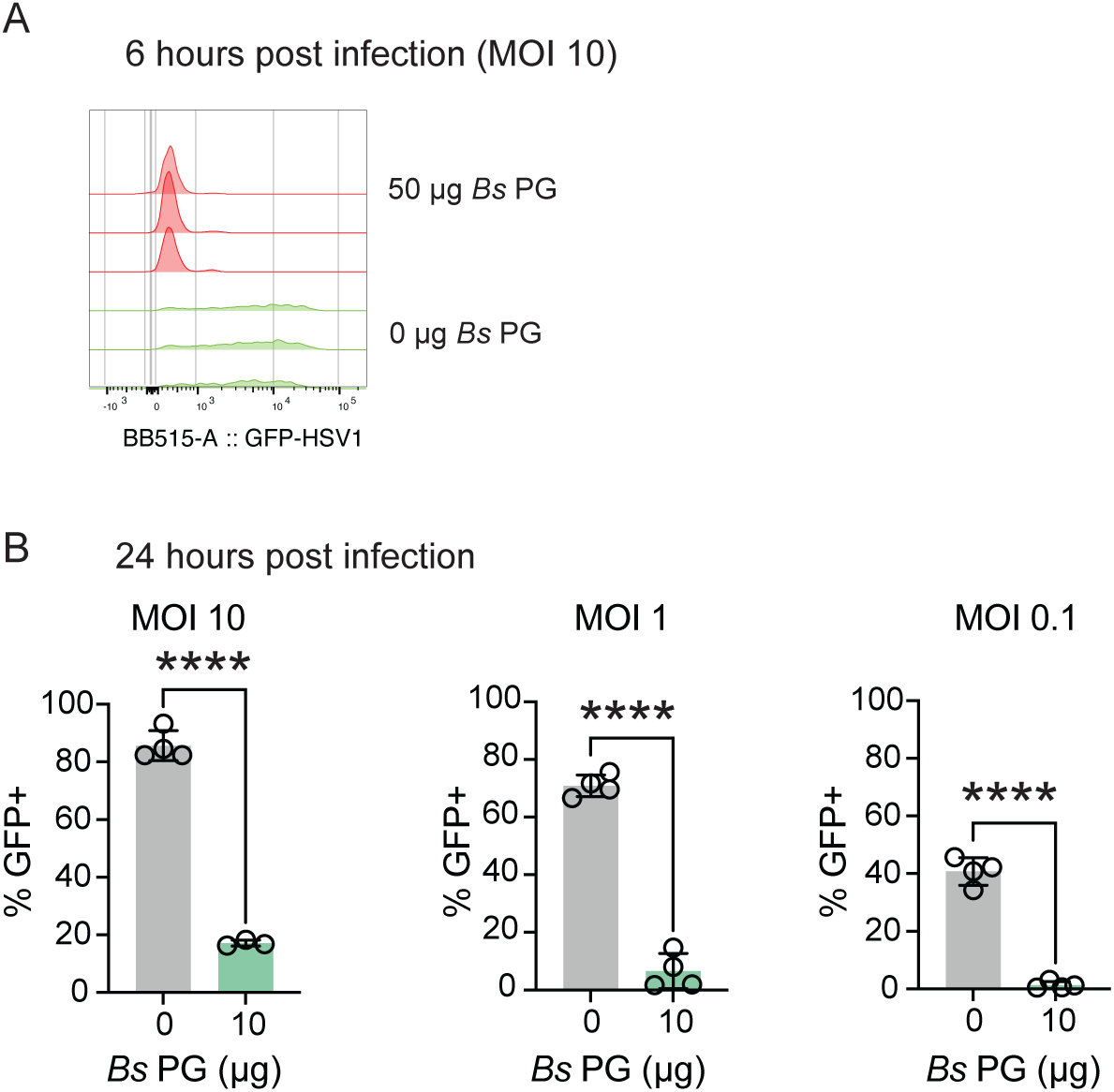
*B. subtilis* PG inhibits HSV-1 infection in HFFs. HFFs were infected at indicated multiplicity of infections with HSV-1 K26GFP strain and indicated amounts of *B. subtilis* PG (Sigma-Aldrich) (A-B). Flow cytometry histograms are shown 6 hours post-infection (A) and frequency of live HSV-1 K26GFP+ infected cells quantified at 24 hours post-infection (B) (n=3-4, a representative experiment is shown). Datasets were compared using unpaired t-tests. Error bars represent the mean with SD.

**Supplementary Figure 6:**
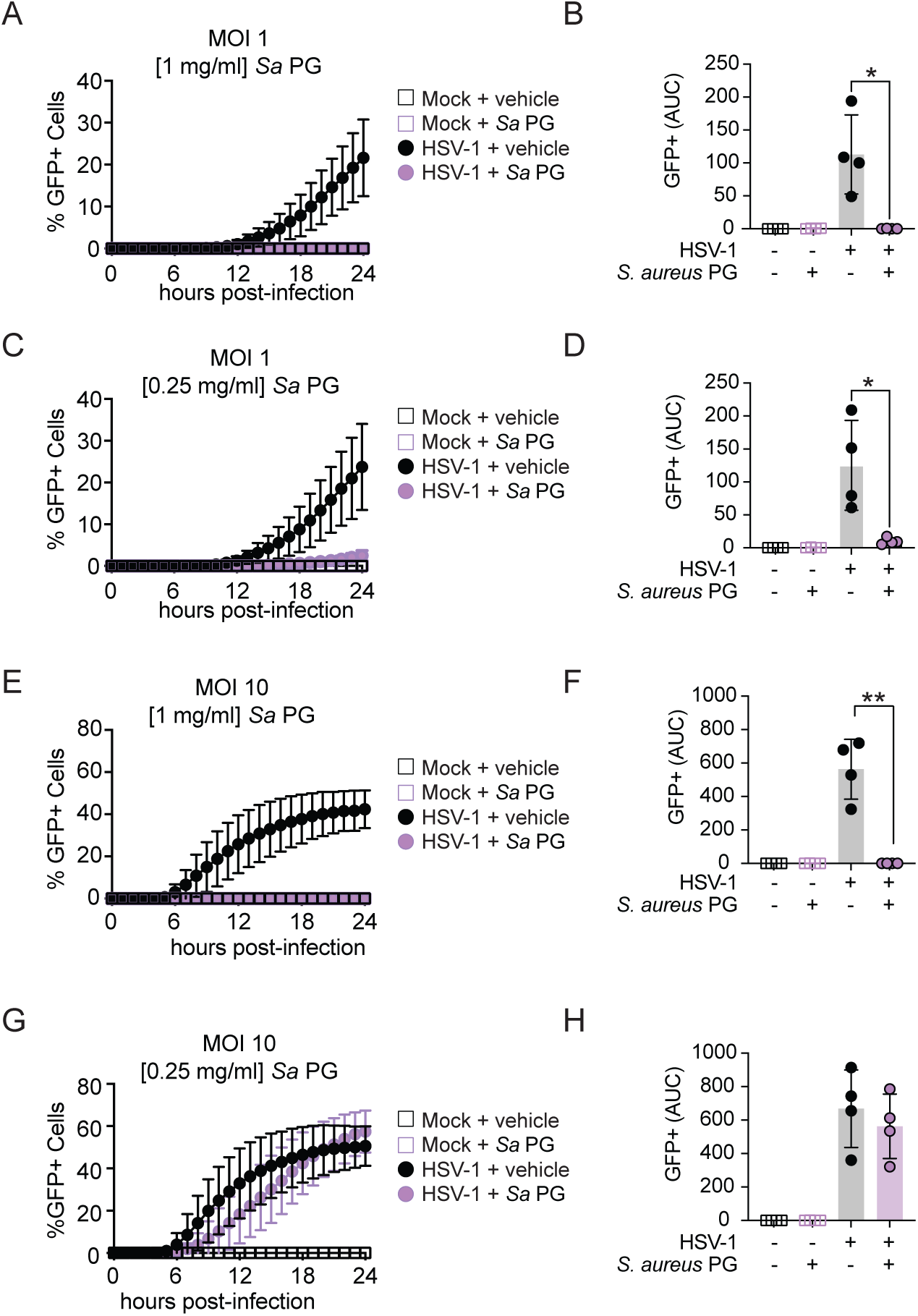
*S. aureus* PG inhibits HSV-1 infection in primary human keratinocytes. Normal human epidermal keratinocytes (NHEK cells) were infected at MOI of 1 (A-D) or MOI of 10 (E-H) with HSV-1 K26GFP and either [1 mg/ml] (A-B and E-F) or [0.25 mg/ml] (C-D and G-H) of *S. aureus* PG (InvivoGen) (n=4 biological replicates across 4 independent experiments). Cells were imaged hourly for 24 hours and ratios of GFP+ to Hoechst+ cells plotted as a measurement of the percentage of HSV-1 K26GFP-infected cells (A, C, E, and G). The area under the curve was quantified (B, D, F, and H) and compared using Welch’s t-tests. Error bars are mean with SD.

**Supplementary Figure 7:**
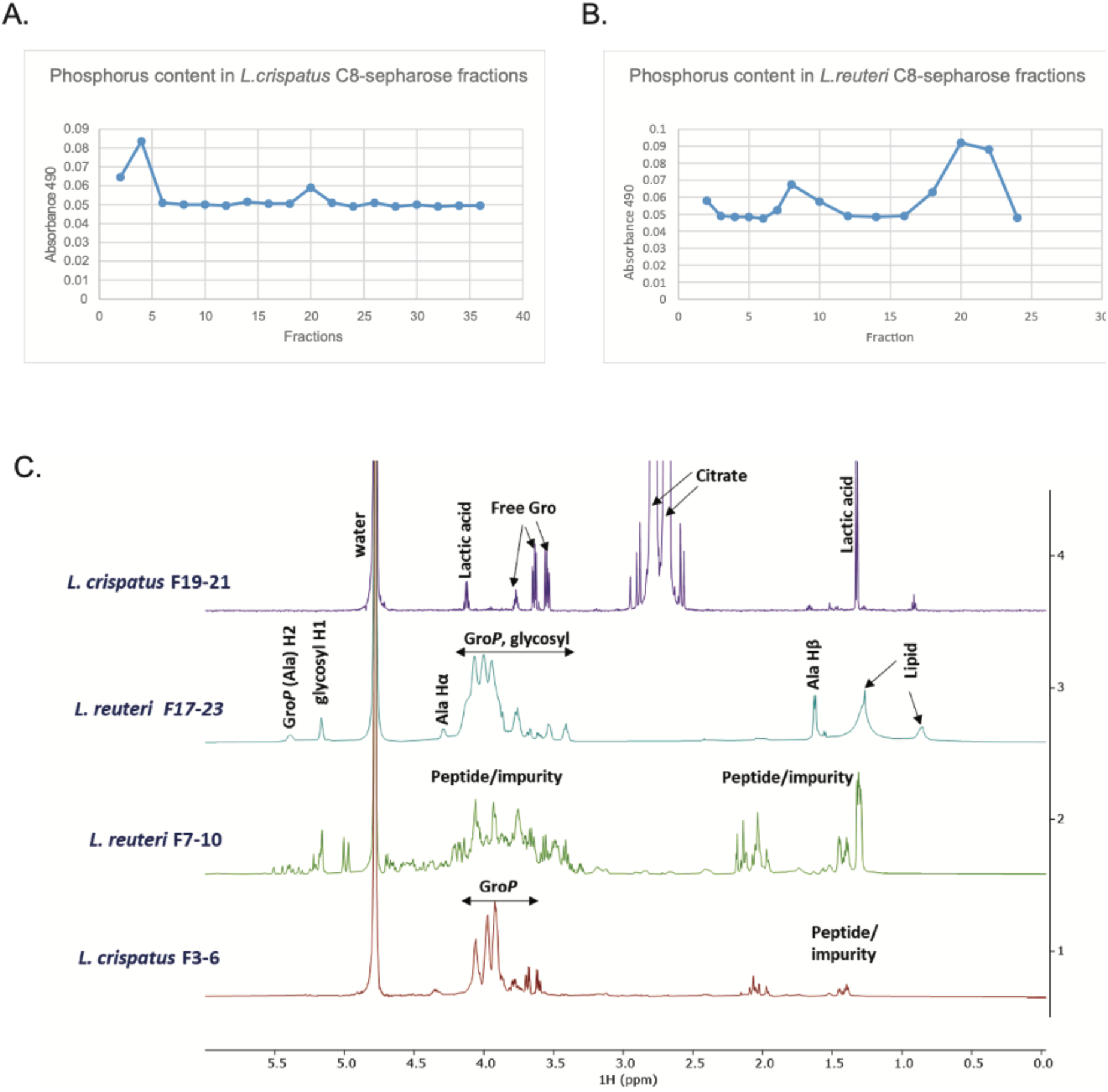
Isolation and NMR purity testing of *L. crispatus and L. reuteri* LTA. (A-B) Absorbance readings of phosphorus assay for C8-sepharose fractions for *L. crispatus* (A) and *L. reuteri* (B) LTA preparations. (C) 1D ^1^H NMR on *L. crispatus* fractions 3-6 and 19-21, and *L. reuteri* fractions 7-10 and 17-23, collected and pooled from the C_8_-sepharose column. The *L. crispatus* F19-21 fractions only contain free glycerol, lactic acid, and contaminating citrate. Fraction F3-6 from *L. crispatus* contains signals around 3.5-4.0 ppm that resemble those of glycerol phosphate (Gro*P*) from LTA, consistent with the presence of a teichoic acid-like polymer. However, there are no or negligible peaks characteristic for lipids around 0.8 and 1.3 ppm, suggesting that this fraction does not contain LTAs. The two LTA fractions extracted from *L. reuteri* differ from *L. crispatus*. *L. reuteri* fraction F7-10 does not show peaks resembling a typical LTA; rather, the peaks are consistent with peptides and other impurities. However, Fraction F17-23 has peaks characteristic for an LTA, as evident from the signals of lipid, substituted glycerol phosphate (Gro*P*), glycosyl residue and alanine, similar to ^1^H signals reported for *S. aureus* LTA (Morath et al., 2001). ^1^H NMR of the material isolated from *L. reuteri* (F17-23) is thus consistent with a pure LTA that is partially alanylated and glycosylated.

**Supplementary Figure 8:**
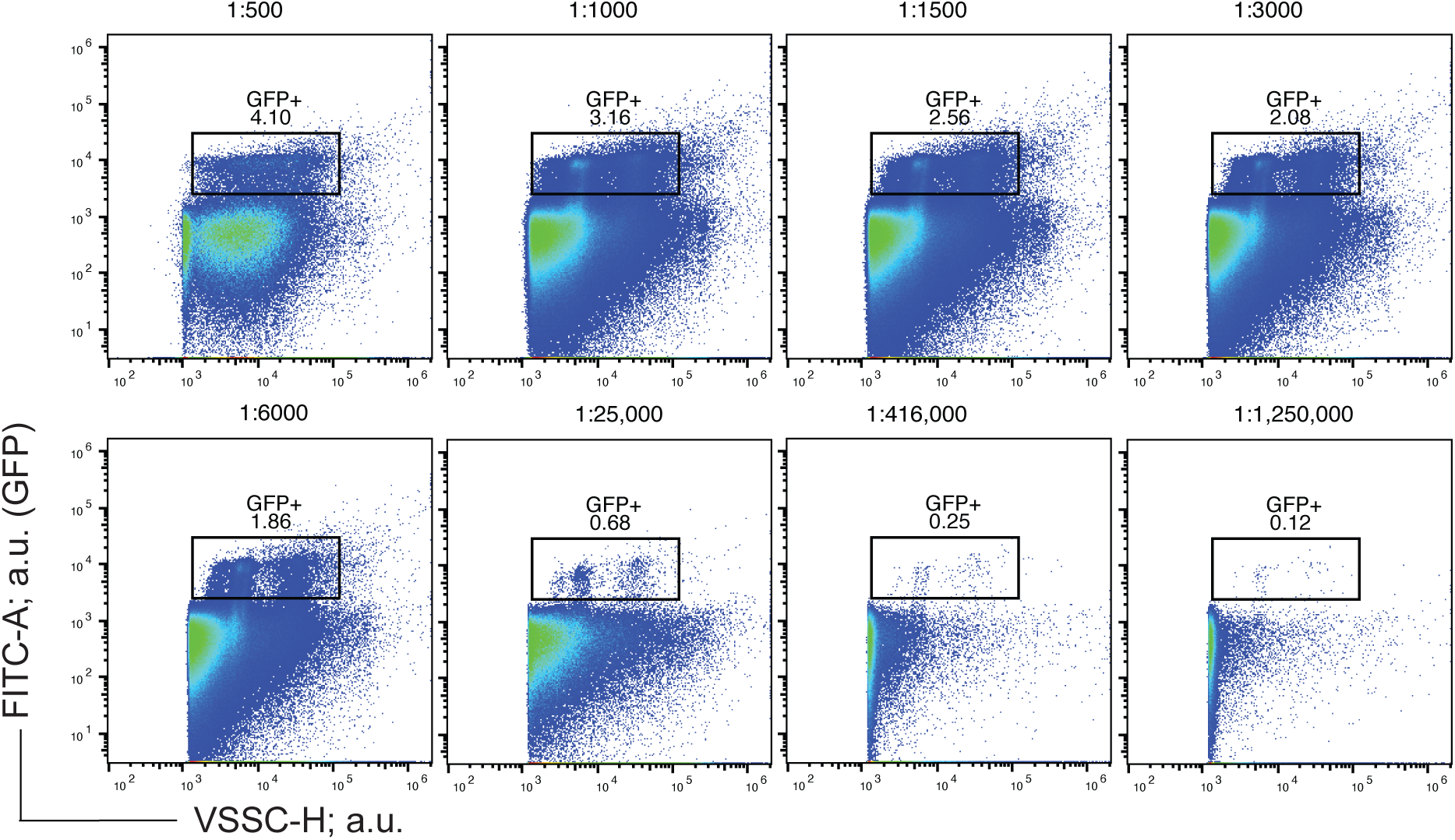
Titrating HSV-1 K26GFP for flow virometry. Serial dilutions of HSV-1 were run to identify the optimal concentrations for acquisition of virions in flow virometry experiments (related to Figure 6). For optimized flow virometry experiments, viruses were diluted ∼25,000 fold from the stock concentration to minimize the chances of coincidental detection. GFP+ virions were gated as indicated on the 1:25,000 plot to indicate the sample concentration used for subsequent flow virometry analysis.

## Notes

### Competing Interest Statement

A.N.D.A and S.G. are co-inventors on a patent related to this work.

### Summary of Updates

Author's name was incorrect and has now been updated.

